# Genomic analysis of the four ecologically distinct cactus host populations of *Drosophila mojavensis*

**DOI:** 10.1101/530154

**Authors:** Carson W. Allan, Luciano M. Matzkin

## Abstract

**Background:** Relationships between an organism and its environment can be fundamental in the understanding how populations change over time and species arise. Local ecological conditions can shape variation at multiple levels, among these are the evolutionary history and trajectories of coding genes. This study examines the rate of molecular evolution at protein-coding genes throughout the genome in response to host adaptation in the cactophilic *Drosophila mojavensis.* These insects are intimately associated with cactus necroses, developing as larvae and feeding as adults in these necrotic tissues. *Drosophila mojavensis* is composed of four isolated populations across the deserts of western North America and each population has adapted to utilize different cacti that are chemically, nutritionally, and structurally distinct.

**Results:** High coverage Illumina sequencing was performed on three previously unsequenced populations of *D. mojavensis.* Genomes were assembled using the previously sequenced genome of *D. mojavensis* from Santa Catalina Island (USA) as a template. Protein coding genes were aligned across all four populations and rates of protein evolution were determined for all loci using a several approaches.

**Conclusions:** Loci that exhibited elevated rates of molecular evolution tended to be shorter, have fewer exons, low expression, be transcriptionally responsive to cactus host use and have fixed expression differences across the four cactus host populations. Fast evolving genes were involved with metabolism, detoxification, chemosensory reception, reproduction and behavior. Results of this study gives insight into the process and the genomic consequences of local ecological adaptation.

## Background

Increasing availability of whole-genome sequencing data provides new insights into the complex relationship between an organism and its environment. By examining changes in the genetic code both at the level of individual genes and at the whole-genome level it is possible to gain a better understanding of how local ecological conditions can shape the pattern of variation within and between ecologically distinct populations [1, 2]. A comprehensive integrative approach combining genomic, phenotypic and fitness data has been identified as the gold standard in understanding the adaptation process [3, 4]. Yet, an examination of the genomic divergence of ecologically distinct populations can yield valuable insight into the adaptation process especially when the genomic data is placed in an ecological context [5]. This later approach can identify genomic regions and loci that exhibit a pattern of variation and evolution suggesting their role in local ecological adaptation. Furthermore, a consequence of the fixation of ecologically-relevant variants has been implicated in the evolution of barriers to gene flow and potentially the origins of reproductively isolated populations, i.e. species [6, 7].

While it has long been accepted that natural selection is a primary driver of change within species as a response to environmental pressures, understanding the mechanism of how this selection leads to speciation is unclear [8, 9]. More recently the idea of ecological speciation, where various mechanisms work to prevent gene flow between populations causing reproductive isolation and eventually speciation, has more directly shown how selection to local ecological conditions may affect the process of speciation [6, 7]. Reproductive isolation interrupts gene flow between populations and may potentially lead to the formation of new species [10]. When different populations of a species inhabits and/or utilizes distinct resources this opens many possibilities for local differentiation that can lead to obstacles of gene flow as these populations are likely to have differing environmental pressures [6, 7]. For example, in the leaf beetle *Neochlamisus bebbianae*, different populations have distinct host preferences and larvae perform significantly worse when growing on alternative host species [8]. Host preferences and performance in this system facilitates the genetic and genomic isolation observed between the host populations, as each prefers a different microenvironment and likely does not interact and hybridize with members of the other population [11, 12].

Comparative genomic studies in mammals have shown clear evidence of positive selection both between humans, mice, and chimpanzees as well as between human populations [13–16]. Genes involved in the immune system, gamete development, sensory perception, metabolism, cell motility, and genes involved with cancer were those found to have signatures of positive selection. While in *Drosophila*, a genome level analysis of 12 species provided insight into the evolution of an ecological, morphological, physiological and behaviorally diverse genus [17]. Findings were relatively consistent with previously studies in other taxa with genes involving defense, chemosensory perception, and metabolism shown to be under positive selection [6, 13, 16, 18]. Since the *Drosophila* 12 genome project [17], several population genomics studies in *D. melanogaster* have examined variation within a single population, between clinal populations and between ancestral (African) and cosmopolitan populations to assess the consequence of population subdivision, evolution of quantitative trait variation and the adaptation to local ecological conditions [19–24]. These genome level analysis have been extended to other *D. melanogaster* species group flies with distinct life history and ecological strategies such as the *Morinda citrifolia* specialist *D. sechellia* [25] and the invasive agricultural pest *D. suzukii* [26].

Studying the sequence level constraints as well as functional categories and networks associated with genes under positive selection is paramount to understanding the process of evolutionary change. However, it is crucial to place patterns of variation and divergence in an ecological context to have a more complete view how selection shapes variation within and between populations. In this study we explore the link between ecology and patterns of genome-wide sequence variation in *D. mojavensis*, a fly endemic to the southwestern United States and northwestern Mexico that has become a model for the understanding of the genetics of adaptation [27]. This species of *Drosophila* is a cactophile in that both larval and adult stages reside and feed in necrotic cactus tissues [28]. *Drosophila mojavensis* has four distinct host populations that are geographically separated (Fig. 1). In addition to geographic separation each population lives on a distinct cactus host species. The four populations are: Santa Catalina Island living on prickly pear cactus *(Opuntia littoralis)*, Mojave Desert living on barrel cactus *(Ferocactus cylindraceus)*, Baja California living on agria cactus *(Stenocereus gummosus)*, and Sonoran Desert living on organpipe cactus (S. *thurberi). Drosophila mojavensis* diverged from its sister species *D. arizonae*, a cactus generalist, approximately half a million years ago [29–32] with the divergence between *D. mojavensis* populations being more recent (230,000 to 270,000 years ago) [33]. Differing host species provide different local environments for each *D. mojavensis* populations. The necrotic cactus environment in which these flies reside is composed not only of plant tissues, but a number of bacteria and yeast species [34–37]. In addition to nutritional differences between the necrotic cactus host, several of the compounds found therein have toxic properties [38–40]. This selective pressure has resulted in the fixation of variants that facilitate the survival of *D. mojavensis* and other cactophilic *Drosophila* species to their local necrotic cactus environment [28, 41].

**Fig. 1.**
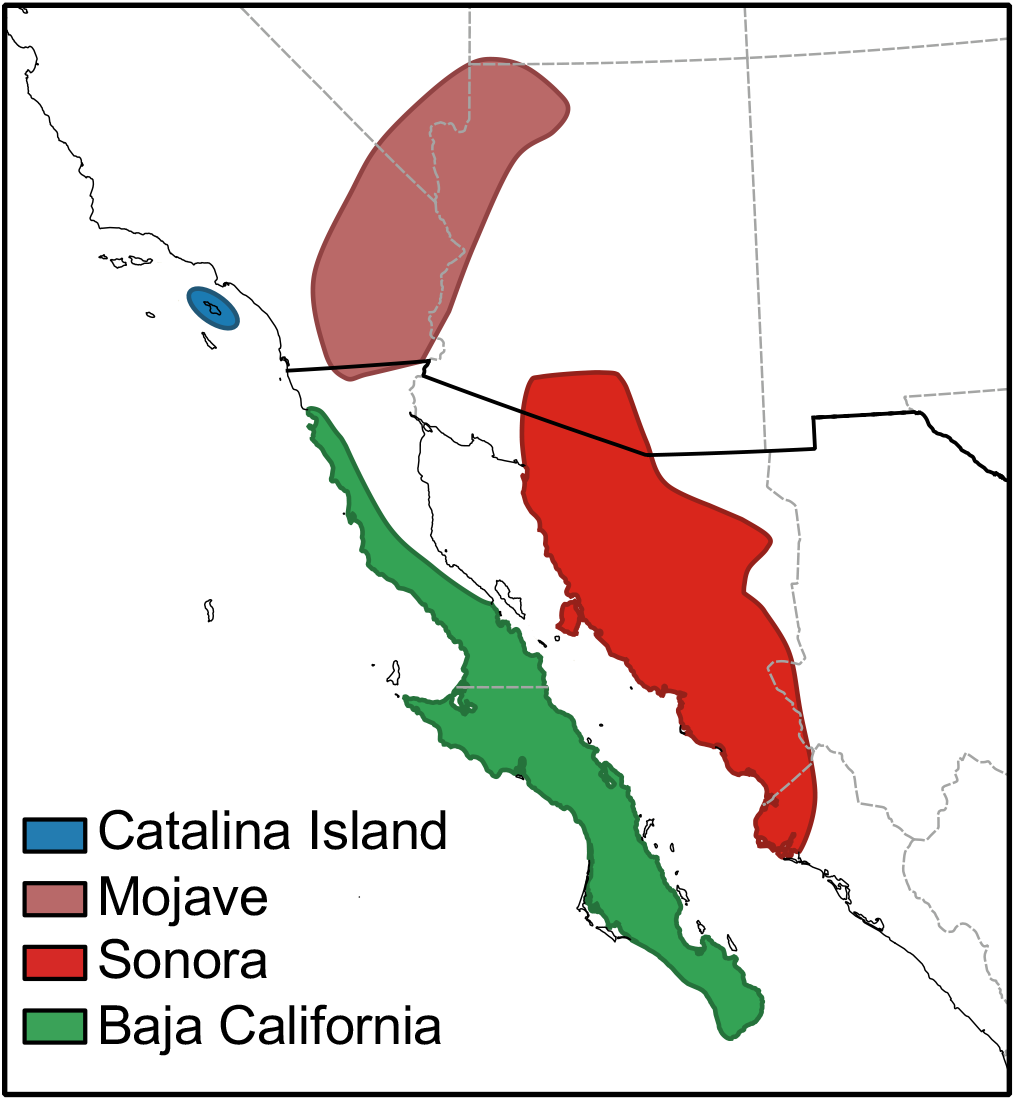
Distribution of the four cactus host populations of *D. mojavensis.*

Population genetics on individual candidate host adaptation genes in *D. mojavensis* has shown evidence for positive selection in loci involved with xenobiotic metabolism [31]. In addition, transcriptome-wide differences have been observed in *D. mojavensis* in response to host shifts [42, 43] as well as indicating fixed expression differences between the host populations [44]. Among the loci that are differentially expressed or constitutively fixed between populations many are involved in detoxification, metabolism, chemosensory perception and behavior, supporting the role of the local necrotic cactus conditions in shaping transcriptional variation [42–44]. Taking into consideration the breadth of ecological information of *D. mojavensis* this study highlights how selection pressures caused by local ecological environments differentially shape patterns of genomic variation across the host populations and provides further insight into how selection acts on organisms and its genome level consequences.

## Results

Number of cleaned reads and the number assembled to the Catalina Island reference genome are shown in Table 1. All three populations had approximately 88 percent of paired-end reads successfully assembled. Mate pair reads had lower rates of mapping ranging from 27 percent to 63 percent. Of the 14,680 loci annotated in the reference genome the vast majority were also present in our template-based assemblies of the other three populations. Of these annotations, a common set of 12,695 were initially processed that did not lack any premature stop codons. From this common set of loci we filtered out those that among the four populations exhibited either less than five total, zero nonsynonymous, or zero synonymous substitutions. This yielded a working set of 9,087 loci for which all subsequent analyzes were performed. The list of all loci examined, summary data, test statistics, and *D. melanogaster* ortholog information can be found in Additional file 1: Table S1.

**Table 1.** Number of cleaned reads and assembled reads for each population.

| Population | Reads Mapped | Total Reads | Proportion Mapped |
| --- | --- | --- | --- |
| Baja California |  |  |  |
| ME | 12,052,662 | 44,912,130 | 0.27 |
| PE | 88,976,029 | 100,263,663 | 0.89 |
| Total | 101,028,691 | 145,175,793 | 0.70 |
| Mojave |  |  |  |
| ME | 26,638,794 | 52,910,406 | 0.50 |
| PE | 73,196,313 | 83,000,942 | 0.88 |
| Total | 99,835,107 | 135,911,348 | 0.73 |
| Sonora |  |  |  |
| ME | 39,962,094 | 63,240,890 | 0.63 |
| PE | 93,857,309 | 105,723,406 | 0.89 |
| Total | 133,819,403 | 168,964,296 | 0.79 |

ME mate pair end reads; PE paired end reads

### Characteristics and patterns of divergence of *D. mojavensis* loci

Estimates of ω (K_a_/K_s_) were calculated using both KaKs Calculator [45] and codeml in PAML [46]. Given that the ω values were highly correlated (r^2^ = 0.88, *P* < 0.001; see Additional file 2: Figure S1) all subsequent analyses were performed using the values obtained from codeml. The distribution of log2 transformed ω are shown in Figure S2. Overall a total of 190 loci exhibited ω values greater than one. When examined per chromosome (Muller Element), we observed that the dot chromosome (Muller F) had the greatest mean ω, followed by the chromosomes for which segregate chromosomal inversions (Muller B and E) and than those chromosomes that lack inversions (Muller A, C and D) (Fig. 2, Additional File 2: Table S2).

**Fig. 2.**
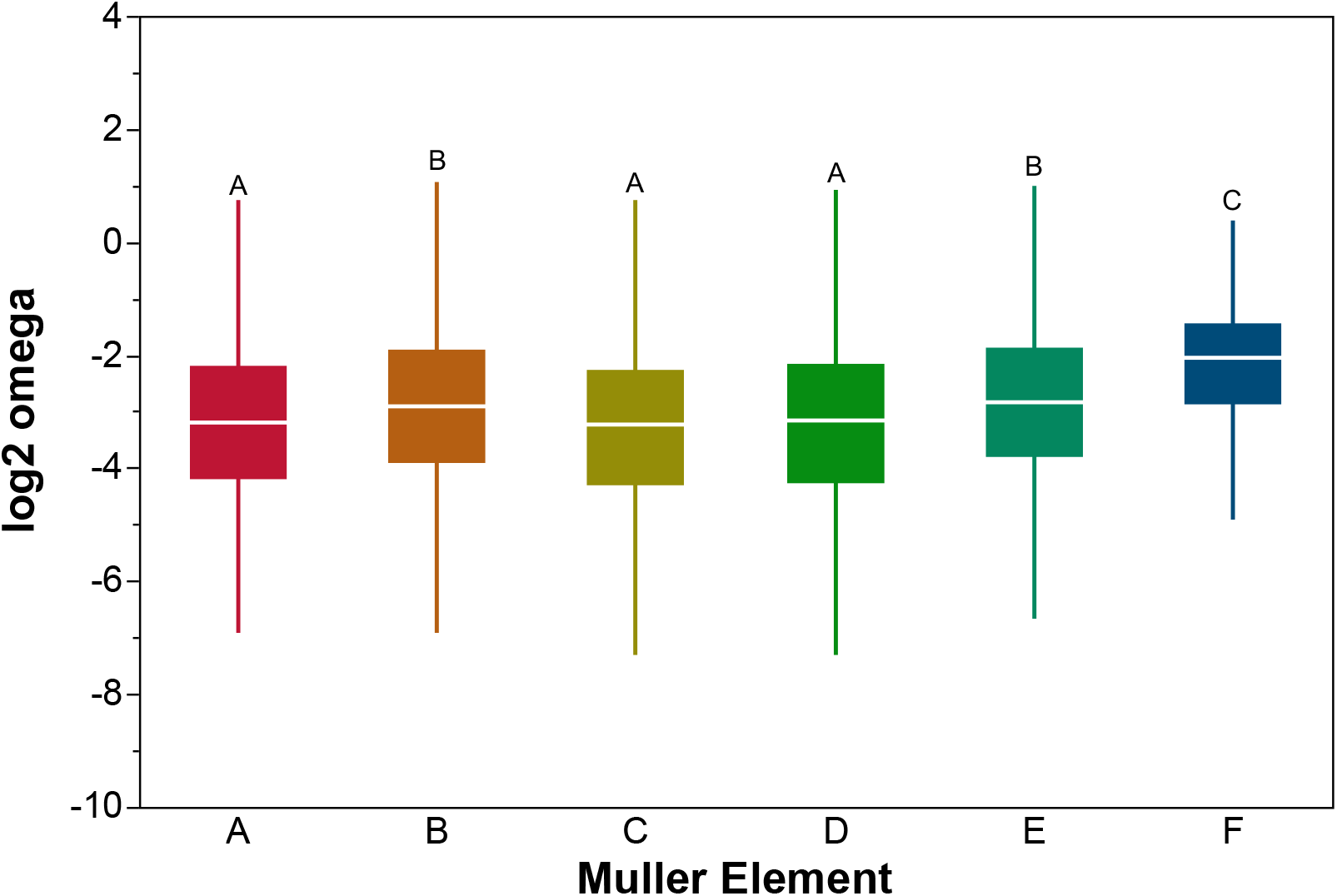
Boxplot of log_2_ ω values for loci located in each of the *D. mojavensis* Muller elements. Elements with different letters are significantly different using a Tukey HSD test (see Table S2).

To describe the characteristics of loci whose evolutionary trajectory could have been shaped by the adaptation of *D. mojavensis* populations to their respective ecological conditions we examined loci with ω values in the top 10% of the distribution, hereafter referred to as TOP10 loci. Furthermore, using codeml we performed a series of gene-wide tests of positive selection for each individual locus. Via a maximum likelihood rate test (model 7 vs. model 8) we identified 912 loci that exhibited a pattern of adaptive protein evolution. We used a smaller set of 244 loci, following an FDR correction, for all subsequent analyses, hereafter referred to as PAML-FDR loci. The set of TOP10, PAML significant loci and those with an FDR correction (PAML-FDR) can be found in Additional file 1: Table S1. The distribution of both the PAML-FDR and TOP10 loci was uniform across the *D. mojavensis* chromosomes (Additional file 2: Figure S3 and S4), with the exception that significantly fewer PAML-FDR genes were present in Muller E (Fisher’s Exact test, *P* = 0.02).

Significant differences in ω values were observed across loci of differing protein coding lengths (Fig. 3). Loci smaller than 1 Kb exhibit significantly higher rate of molecular evolution, followed by those 1-2 Kb and then by gene categories of longer lengths (Additional file 2: Table S3). A similar pattern of ω values was observed for the TOP10 loci, where a significant excess of the smaller gene group (< 1 Kb) was composed of TOP10 loci, and a significantly fewer were observed in the greater than 4 Kb bin (Additional file 2: Figure S5). Although the overall ω was greater in shorter loci, the proportion of these loci who exhibited a significant pattern of positive selection was significantly less (Additional file 2: Figure S6). Similarly to what was observed for gene length, genome-wide, loci with fewer exons tended to have greater levels of ω, with the highest observed from loci having two exons, then those with either only one or three exons, followed by those having four to six exons and lastly those with seven or more (Additional file 2: Figure S7, Table S4). TOP10 loci were overrepresented in the one and two exon categories and underrepresented in the more than seven exon category, whereas the PAML-FDR loci where uniformly distributed across all exon number categories (Additional file 2: Figures S8 and S9).

**Fig. 3.**
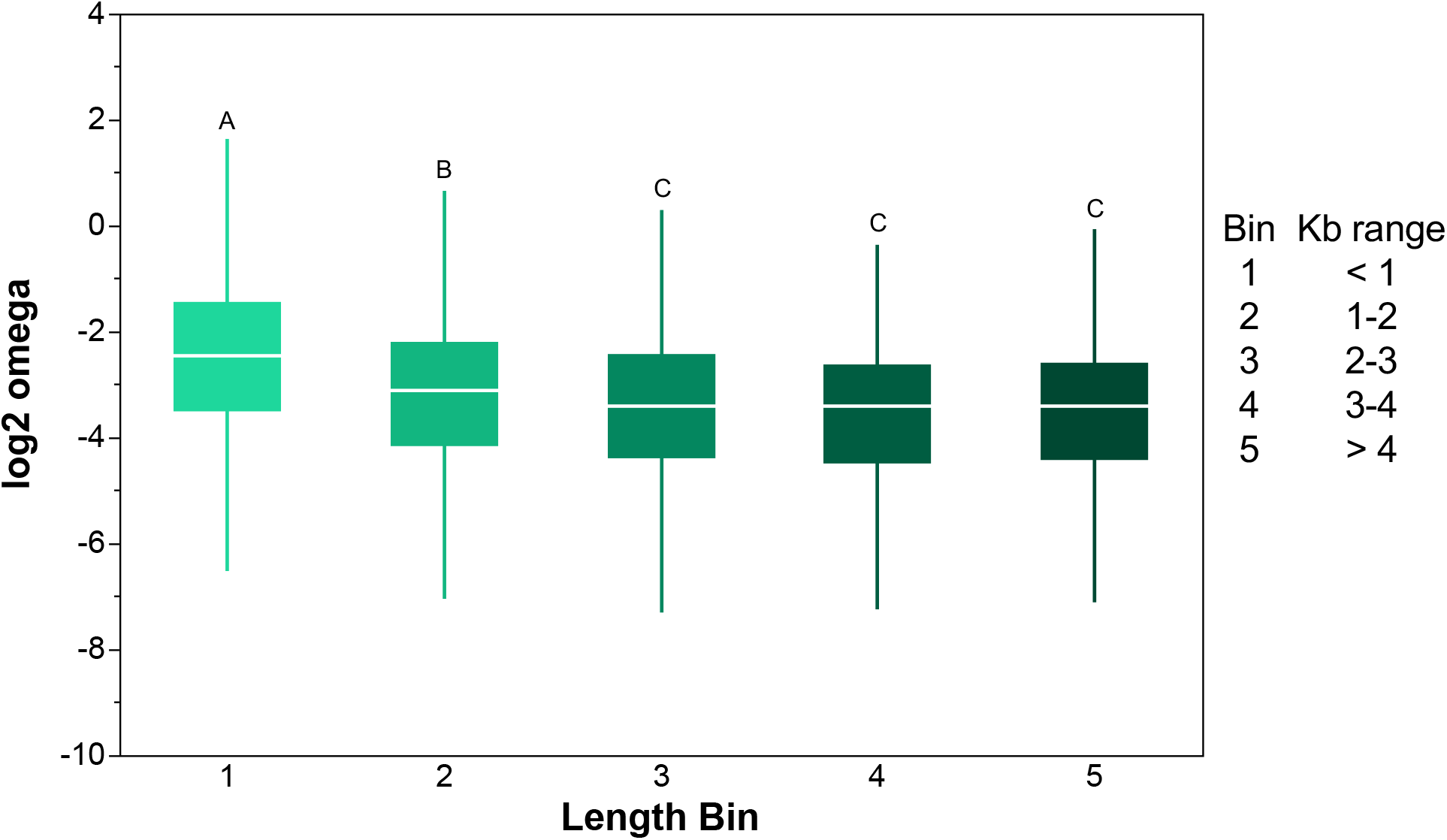
Boxplot of log_2_ ω values for loci in five different coding length bins. Bins with different letters are significantly different using a Tukey HSD test (see Table S3).

### Relationship between expression and rate of molecular evolution

To assess the relationship between expression level and rate of molecular evolution we integrated our results with previous collected RNAseq data from *D. mojavensis* [47]. When examined genome-wide, genes with male-biased expression had significantly greater ω values than female-biased (Tukey HSD, P < 0.001) and unbiased (Tukey HSD, P < 0.001) expressed genes, and female-biased genes had the lowest rate (Tukey HSD, P < 0.001) of molecular evolution of all three expression categories (Additional file 2: Figure S10, Table S5). Among the TOP10 loci, there was a significant representation of them in the male-biased group of genes and a significant underrepresentation in the female-biased genes (Fig. 4). No significant over-or underrepresentation was observed among the PAML-FDR genes with respect to the sex biased expression categories (Additional file 2: Figure S11). Expression data was also used to assess the relationship between overall expression level and rate of molecular evolution. After removing both the female- and male-biased genes, we observed that of the 5,101 remaining loci those in the lowest expression category showed the greatest ω values (Additional file 2: Figure S12, Table S6). Similarly, the TOP10 loci were overrepresented among the low expression category of loci and no differences were observed among the expression categories of the PAML-FDR loci (Additional file 2: Figures S13 and S14).

**Fig. 4.**
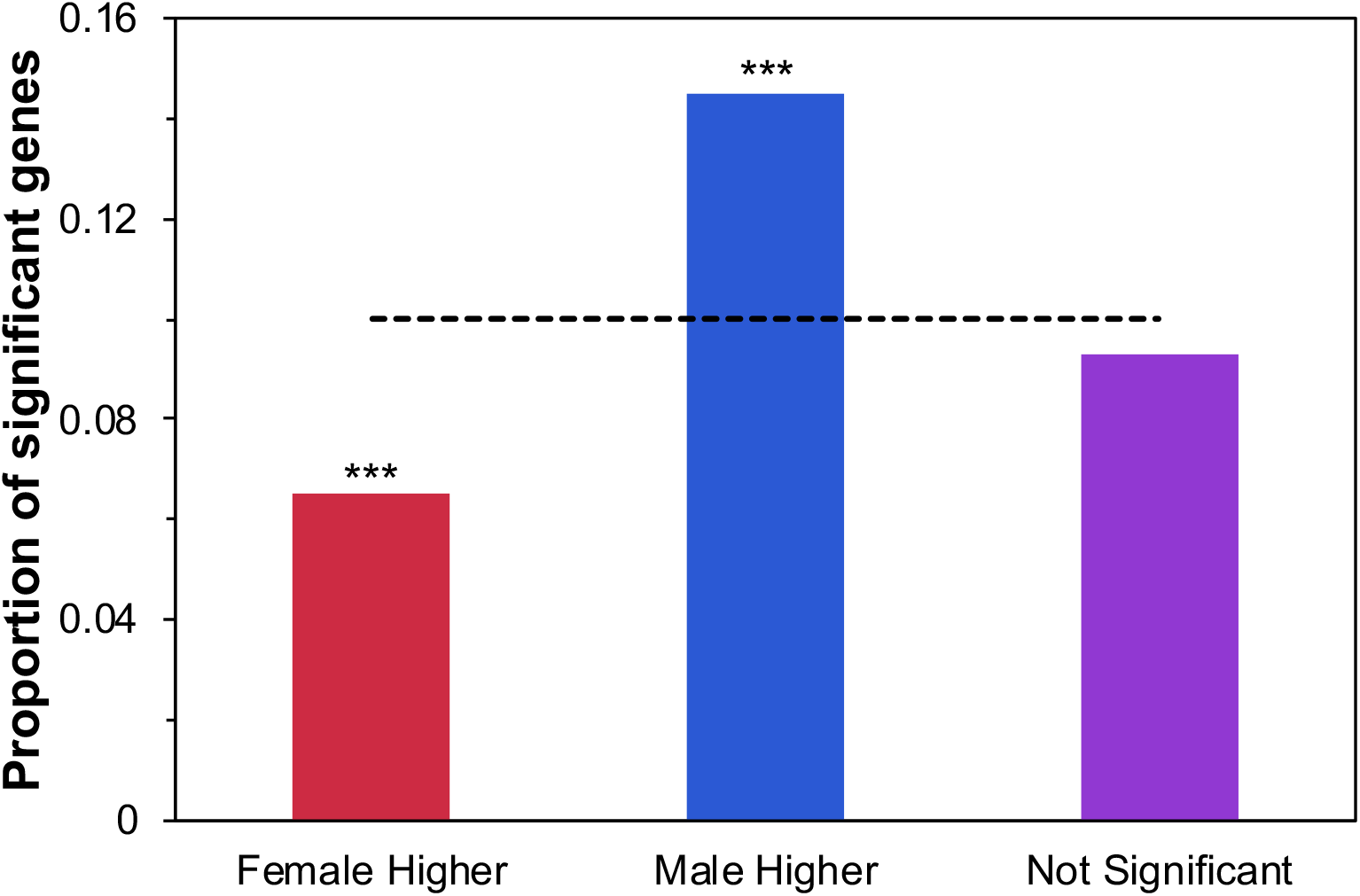
Proportion of TOP10 loci that show female-bias, male-bias or unbiased gene expression. Dashed line indicates the genome wide proportion of TOP10 loci (0.10). Gene expression data is from [47]. Asterisk indicate significance via Fisher’s Exact test (* P < 0.05, ** P < 0.01, *** P < 0.001).

We also integrated our genomic data with two prior ecological transcriptional studies. We compare rates of molecular evolution of loci that are differentially expressed in response to cactus host utilization [43] as well as those loci who exhibit fixed significant expression differences between the four host populations in the absence of cactus compounds (i.e. constitutive differences) [44]. To remove the potential confounding effect of those loci that show a pattern of positive selection, we removed those loci from the subsequent expression analysis. For both datasets, loci that are either differentially expressed in response to necrotic cactus (*P* < 0.001 post FDR correction) or those that show constitutive differences between the populations (*P* < 0.001 post FDR correction) have a significantly greater value of ω (ANOVA, *P* < 0.001, for both comparisons) (Additional file 2: Figures S15, Table S7).

### Functional gene groups analysis

Of our 9,087 genes in our filtered dataset, approximately 14% (1,238) genes did not have orthologous calls back to loci in the *D. melanogaster* reference genome (Additional file 2: Figure S16). Of the remaining set of genes with *D. melanogaster* orthologs, less than half of the genes (3,649) had at least one gene ontology (GO) term. The percentage of loci without *D. melanogaster* orthologous in the TOP10 and PAML-FDR genes was greater (40% and 23%, respectively). Overall only 336 and 144 loci had at least one GO term for the TOP10 and PAML-FDR datasets, respectively. Clustering of biological process and molecular function GO terms within the TOP10 and PAML-FDR dataset illustrated some distinct functional groups. Fig. 5 illustrates the biological process functional clusters for TOP10 genes, in which clusters associated with reproduction/development, detoxification and response to stimuli, and behavior are present. A network analysis of the same set of loci indicates similar functional networks as well as those associated with defense and chromatin regulation and remodeling (Fig. 6). Functional and network clustering for molecular function GO terms, KEGG and the PAML-FDR dataset can be found in Additional file 2: Figures S17-S20, Additional file 3: Table S11. Among molecular functions, in the TOP10 dataset, serine endopeptidase activity appeared to be overrepresented (Additional file 2: Table S8).

**Fig. 5.**
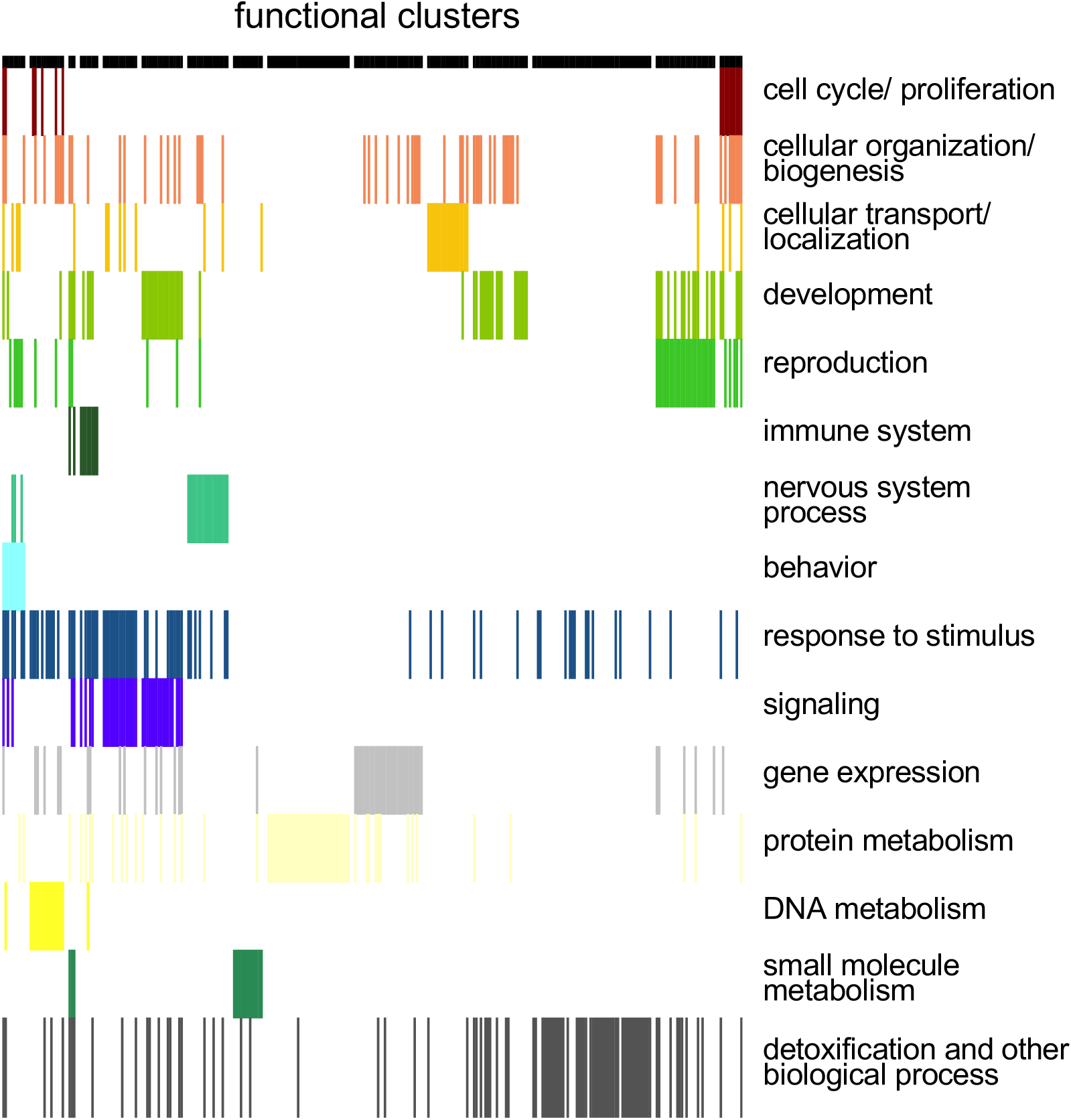
Functional clustering of Biological Process GO terms of the TOP10 loci. Details of gene composition of each cluster is in Additional file 3: Table S11.

**Fig. 6.**
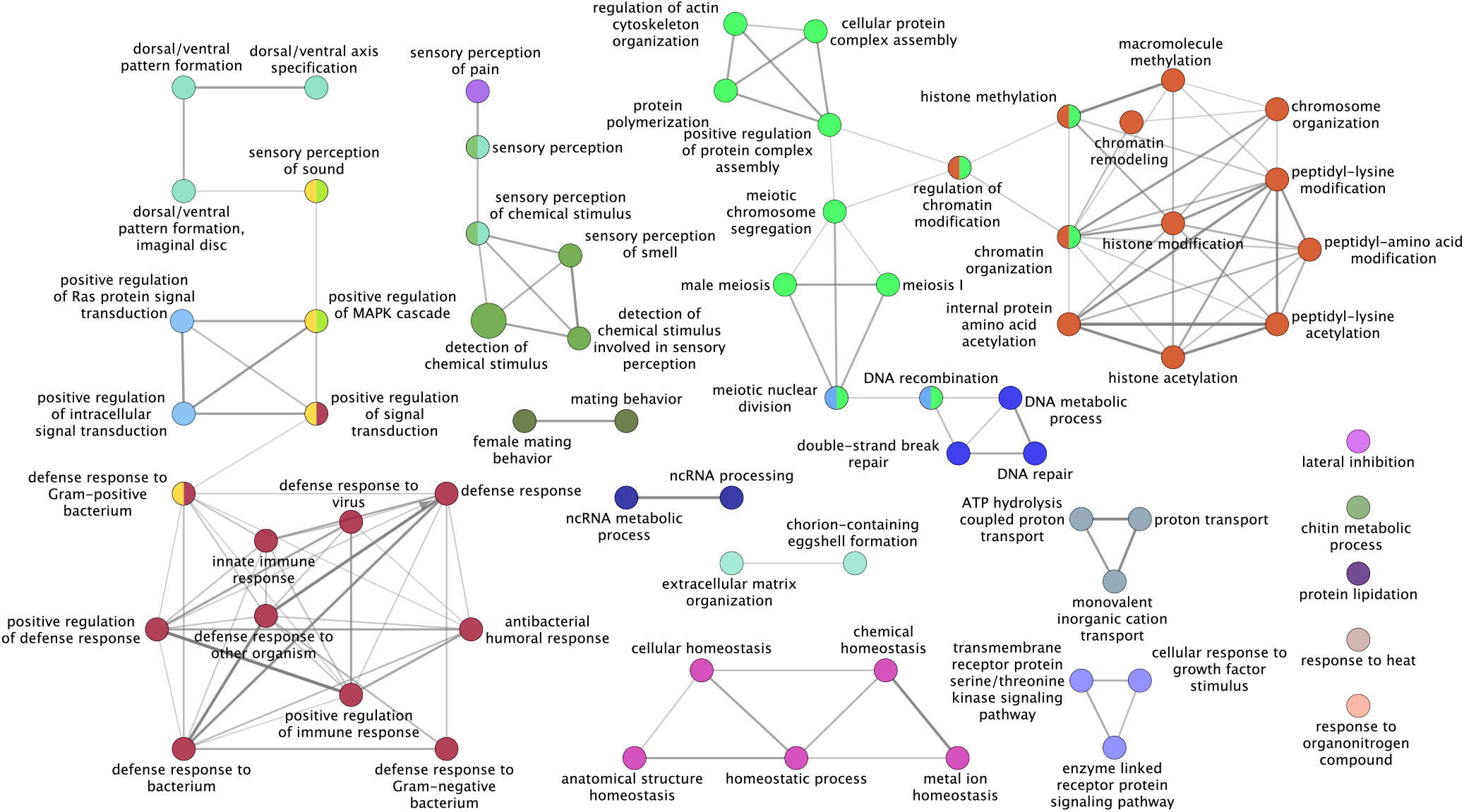
Network clustering of Biological Process GO terms of the TOP10 loci. Network clustering was performed using ClueGo using the following parameters: Min GO Level = 3, Max GO Level = 8, All GO Levels = false, Number of Genes = 3, Get All Genes = false, Min Percentage = 5.0, Get All Percentage = false, GO Fusion = true, GO Group = true, Kappa Score Threshold = 0.3, Over View Term = Smallest PValue, Group By Kappa Statistics = true, Initial Group Size = 1, Sharing Group Percentage = 50.0.

## Discussion

In this study we sequenced, assembled and analyzed the genomes of each of the four cactus host populations of *D. mojavensis* for the purpose of assessing the genomic consequences of the adaptation to local ecological conditions. Overall, we were able to analyze the sequence, pattern of divergence and structure of 9,087 genes. And although the four genomes examined diverged relatively recently [29–33], for several loci, sufficient number of substitutions occurred for us to begin to assess the changes associated with cactus host adaptation.

Unlike what is present in *D. melanogaster, D. mojavensis* chromosomes are all acrocentric and its karyotype is composed of six Muller elements [48]. In *D. melanogaster* element A is the X chromosome and elements B/C and D/E form large metacentric chromosomes (2L/2R and 3L/3R, respectively), while the F element or dot chromosome is reduced in sized and highly heterochromatic [49, 50]. In *D. mojavensis* we observed the highest rate of molecular evolution in the small F element, followed by elements B and E, and then the remaining autosomal elements and the X chromosome (Fig. 2).

Selection on the X chromosome has been examined in a number of studies with somewhat variable results [51]. Analysis of several melanogaster group species has shown significant elevated ω values for genes on the X chromosome [17]. From population genetics theory it is generally predicted that the X chromosome would show elevated rates of evolution due to its reduced population size and level of recombination [51]. A subsequent genomic analysis of the X chromosome across more distant *Drosophila* species *(D. melanogaster, D. pseudoobscura, D. miranda* and *D. yakuba)* failed to find evidence of increased protein evolution on the X chromosome [52]. It is difficult to make any conclusions about the lack of a pattern of accelerated X chromosome evolution found here, it may be possible that there has not been enough divergence time between these populations for factors such as effective population size to have a measurable effect. The greatest ω values were present in the dot chromosome which in *D. mojavensis* is heterochromatic and has a highly reduced level of recombination [53], which would make it highly susceptible to sweeps and hence higher rates of molecular evolution.

Within *D. mojavensis* there are polymorphic inversions in Muller elements B and E [54], both exhibited overall higher chromosomal-wide levels of ω (Fig. 3). Lower levels of recombination and higher divergence rates have been known to occur around the inversion breakpoint regions in *Drosophila* [55]. One possible explanation for the elevated rates of molecular evolution in these chromosomes is the distinct karyotypes of the sequenced lines (Additional file 2: Table S9). One consequence of a template-based assembly as performed in this study, is that chromosomal structural differences can be largely wiped away. A more detailed analysis of the consequence of chromosomal inversion on the evolutionary trajectories of associated loci will be performed in future analyses of *de novo* assemblies of *D. mojavensis* genomes from all host populations as well as from sibling species *(D. arizonae* and *D. navojoa)* (unpublished data, Matzkin).

Genes across the genome as well as those with evidence of positive selection or in the top 10 percent of ω values were assessed for a number of characteristics. Genome-wide loci exhibiting greater ω values tended to be shorter, have fewer exons (3 or less), have low expression, be differentially expressed in response to cactus host use and have fixed expression differences across the four cactus host populations of *D. mojavensis* (Fig. 3; Additional file 2: Figures S7, S12, S15). Overall this pattern of divergence was similar when examining the TOP10 or PAML-FDR loci. Previous genomic analyses in *D. melanogaster* and related species have observed similar characteristics of loci with elevated ω values. This indicates that although the phylogenetic scale of the present study is limited (within *D. mojavensis)* the forces shaping genome evolution between diverged species can also be observed between recently isolated populations within species.

The first comparative genomic study within the *D. melanogaster* group species [56] observed an association between coding length and ω, which they partially attributed to a positive correlation between K_s_ and protein length. Longer genes have more of these mutations and this may explain in part why genes with high ω values are likely to be shorter. In this study we did not observe such correlation, in fact the relationship is negative (P < 0.001), but explains very little of the variation in K_s_ (r^2^ = 0.004) (Additional file 2: Figure S21). Therefore, it is difficult to infer the effect of the association between K_s_ and protein length, and the lack of positive correlation might be a function of the close relationship between the genomes studied here. The negative association between intron number and rate of molecular evolution has been previously suggested to be due to the presence of exonic splice site enhancers which help in the correct removal of introns from the transcription sequence. As mutations in these regions are more likely to be conserved changes here could cause an intron to not be removed or part of an exon to be removed instead [57]. The link between intron presence and ω values may also help explain why TOP10 genes tend to be shorter as long genes are more likely to have introns [58]. The correlation between gene length and rate of molecular evolution could also be explained as a result of the increased level of interactions between sites of larger exons [59]. In this study a negative correlation between ω and exon length (r^2^ = 0.08, *P* < 0.001) was observed (Additional file 2: Figure S22). These interactions between residues of a protein, commonly refer to as Hill-Robertson interference [60], have a tendency to buffer against the accumulation of amino acid substitutions and can explain a significant portion of the pattern of molecular evolution in genomes [61]

Highly expressed genes tend to have a higher level of constraint as indicated by the tendency of having lower rates of molecular evolution. This has been previously explained as being a result of selection against mutations that alter transcriptional and translational efficiency as well as selection for the maintenance of correct folding (translational robustness) [56, 62–66]. Given our coarse transcription data we were not able to tease apart which of the abovementioned forces might more strongly shape the rate of molecular evolution in these genomes. Nonetheless we observed a clear negative relationship across the four *D. mojavensis* genomes between transcriptional level and ω. In addition to overall expression, both tissue and sex-bias expression have been known shape the evolutionary trajectories of genes [61, 67–69]. Male, or more specifically testes expressed genes have been associated with elevated rates of molecular evolution in Drosophila and across many taxa [70]. Many of these loci are believed to be under strong sexual selection, which would explain their accelerated rate of molecular evolution. As predicted we observed an overall higher rate of molecular evolution in male-biased genes. Even female-biased loci exhibited a significant greater ω than unbiased genes. Previous behavioral and molecular studies in *D. mojavensis* have shown that this species experiences strong and recurrent bouts of sexual selection [71–78].

Loci indicating a pattern of positive selection and those with elevated ω appear to be associated with a wide range of metabolic processes. These changes are likely a result of the distinct nutritional and xenobiotic environment the different *D. mojavensis* populations experience. The chemical composition of the cacti and the species of yeast found in each rot varies [34–41] and thus the populations have likely needed to optimize the recognition, avoidance and processing of these necrosis-specific compounds through changes in metabolism, physiology and behavior.

One aspect of metabolism that has likely been shaped by cactus host adaptation is the detoxification of cactus compounds, as the distinct cactus hosts have different chemical compositions. Expression studies have shown that genes involved in detoxification are enriched when flies develop in an alternative necrotic cactus species. Fitness costs of living on the alternative cactus have also been shown to be quite high with those flies having low viability (< 40%) [43, 79, 80]. Out of all GO terms examined in this study, the only ones that were consistently overrepresented were those associated with serine-type endopeptidase activity. These type of proteins perform a number of function within organisms, among them is their targeting of organophosphorus toxins [81]. These compounds are often used in pesticides and are found to inhibit serine hydrolase function in both insects and vertebrates [81]. While the apparent positive selection on these genes could be due to a response to pesticides they might experience in the field, but more likely they may be evolving in response to the effects of the toxic or nutritional compounds found in cactus rots.

Cactophilic Drosophila have been shown to deploy a number of enzymatic strategies to ameliorate the deleterious consequences of ingesting cactus necrosis-derived compounds. Many of the previously identified proteins playing a role in detoxification in cactophiles (Glutathione S-transferases, Cytochrome P450s, Esterases and UDP-glycosyltransferase) have been associated with detoxification in a broad number of taxa [82–86]. In fact, in recent comparative genomic analysis of the cactophilic *D. buzzatii* [87] and *D. aldrichi* [88], a number of metabolic genes, including those associated with detoxification were shown to be under positive selection. In the present genomic analysis of the *D. mojavensis* genome we observed that the largest functional cluster (Fig. 5) was composed of several genes belonging to known detoxification protein families, such as Cytochrome P450 and Glutathione S-transferases (Gst). Furthermore, previous transcriptional studies have indicated that these same categories of detoxification loci are differentially expressed when *D. mojavensis* are utilizing necrotic cactus tissues [42, 43]. A population genetics analysis of *GstD1* has indicated a pattern of adaptive amino acid evolution at this locus in the Sonora and Baja California populations [31]. The location of the fixed residue fixed in the lineages leading to these two populations indicated potential functional consequences and a recent kinetic analysis of these proteins have support this prediction (Matzkin, unpublished data).

The diversity of bacterial species found on each necrotic cactus provides, directly or indirectly, nutritional resources for the fly populations, but also are composed of potentially distinct pathogenic organisms [89, 90]. A number of genes with elevated rates of molecular evolution in this study are linked to a range of processes involved with the immune response. As each population is faced with a different composition of threats, the evolutionary arms race between flies and their pathogens creates further divergence between the populations as they face different pathogenic landscapes. Studies in other species, such as *D. simulans*, have found that genes with immune related functions were found to have higher rates of positive selection than the genome average [91]. Exposure to bacterial pathogens in *D. mojavensis* could occur while utilizing the necrotic cactus substrate, but as has been previously suggested [92], via sexual transmission.

A number of the TOP10 loci in this study perform functions associated with sensory perception and behavior (Fig. 6). *Drosophila mojavensis* larvae actively seek out patches of preferred yeast species [93] and across the four host populations there are distinct larval foraging strategies [94]. More specifically genes involved in chemosensory behavior were observed to have elevated ω values in these genomes. Across Drosophilids, there have been a number of studies indicating the links between the evolution of chemosensory genes and host specialization [95–97]. In *D. sechellia*, a specialist species, was found to be losing olfactory receptor genes at a faster rate than its sibling generalist species *D. simulans* [98]. In *D. mojavensis* each cactus species rot contains different compounds and thus have distinct set of volatiles emanating from the necroses [39, 40]. These chemical differences have shaped the feeding and oviposition behavior of flies as has been shown by the exposure of adults to cactus volatiles [99–101]. Recent analysis of populations differentiation in odorant and gustatory receptors have shown that unlike what might be initially predicted a number of the changes in these receptors suggests that effects at the level of signal transduction in addition to odorant recognition [102]. Further functional analysis is needed to better understand the evolution and functional changes of chemosensory pathways associated with the adaptation to necrotic cacti.

In addition to their role in xenobiotic metabolism, serine proteases have been shown to be involved in the network of proteins associated with reproductive interactions in several taxa. In *D. melanogaster* accessory gland proteins (ACP), such as sex peptide, are found to perform a wide range of functions ranging from stimulating ovulation and reducing a female’s remating rate to helping to defend against infections [103–105]. Knockouts of serine proteases have been shown to interfere with the behavioral and physiological effects of the male-derived sex peptide [105]. In *D. mojavensis* and its sister species *D. arizonae* a large number of proteases are expressed in female reproductive tracts and several have been shown to be under strong positive selection [74, 106–108]. In addition to ACPs being transferred via the ejaculate, gene transcripts have been found to be deposited by males into females during copulation [73]. Some of these male-derived transcripts could alter the female’s transcriptional response, while other may potentially be translated within females. Furthermore, the loci of several of these male-transferred transcripts show a pattern of strong and continuous positive selection, likely as the result of persistent sexual selection [72]. While there seems to be no postzygotic effects of sexual isolation within the *D. mojavensis* populations there is some evidence of prezygotic isolation, where certain populations prefers to mate with members of its own population [77]. The pattern of positive selection and/or elevated rate of molecular evolution for proteases and reproductive loci in the present study may highlight the continuing genomic consequence of sexual selection in this species.

## Conclusions

Local ecological adaptation can shape the pattern variation at multiple levels (life history, behavior and physiological), and the imprint of this multifaceted selection can be observed at the genomic level. In this first ever genome-wide analysis of the pattern of molecular evolution across the four ecologically distinct populations of *D. mojavensis*, we have begun to describe the genomic consequences of the adaptation of these cactophilic *Drosophila* to their respective environments. Given that across the four populations are known differences in cactus host use, which encompass differences in both toxic and nutritional compounds, but as well as necrotic host density, temperature, exposure to desiccation and likely pathogens and predators, it was expected that a number of functional classes of loci might be under selection. Among genes with elevated rates of change are those involved in detoxification, metabolism, chemosensory perception, immunity, behavior and reproduction. We observed general patterns of variation across the genomes indicating that loci with elevated rates of molecular evolution tended to be shorter, with fewer exons and have low overall expression. Furthermore, fast evolving loci also were more likely to be differentially expressed in response to cactus host use and have fixed inter-population expression differences, indicating that both transcriptional and coding sequence changes have been involved in the local ecological adaptation of *D. mojavensis.*

## Methods

### *Drosophila mojavensis* lines and sample preparation

Fly lines MJBC 155 collected in La Paz, Baja California in February 2001, MJ 122 collected in Guaymas, Sonora in 1998, and MJANZA 402-8 collected in ANZA-Borrego Park, California in April 2002 were used as the source lines for the sequencing of three *D. mojavensis* populations. These lines were highly inbred to reduce the heterozygosity of their DNA. Summary of the karyotype of each of the lines sequenced as well as the Catalina Island template genome stock (15081-1352.00) can be found in Additional file 2: Table S9. The flies were grown for two generations in banana molasses media [94] supplemented with ampicillin (125 μg/ml) and tetracycline (12.5 μg/ml), to prevent the isolation of bacterial DNA in addition to the flies’. DNA was extracted from homogenized whole male flies using a combination of phenol/chloroform DNA extraction and Qiagen DNeasy spin-columns to achieve the required amount of DNA material. RNase A was used to reduce RNA contamination. Gel electrophoresis was run on each sample to check the quality of the extraction. Any samples with RNA contamination were run through a Qiagen QIAquick PCR Purification Kit spin column to filter contaminates. Extracted DNA was sent to the HudsonAlpha Institute for Biotechnology Genomic Services Lab (Huntsville, Alabama) for sequencing. One hundred base pair paired-end and mate pair sequencing was done on an Illumina HiSeq 2000 with one lane for each.

### Genome assembly

Paired-end and mate pair Illumina reads were filtered and trimmed using step one of the A5 Pipeline [109]. This step uses SGA [110] and TagDust [111] with the quality scores from the Illumina FASTQ files to reduce the number of low quality reads. A5 was run on the Dense Memory Cluster of the Alabama Super Computer Center with four processing cores and 64 gigabytes of memory allocated for each run. With the reads cleaned they were assembled to the template genome. The reference genome of the Catalina Island population of *D. mojavensis* was assembled as part of the *Drosophila* 12 Genomes Consortium [17]. Version 1.04 of the reference genome was retrieved from FlyBase version FB2015_02 [112]. From the reference sequence, genome scaffolds [113] containing the protein-coding genes previously mapped to a chromosome, were extracted for use as a template for the assembly; these scaffolds are detailed in Additional file 2: Table S10. The reference templates as well as the Illumina reads were imported into Geneious 8.1. Assembly was done separately for paired-end and mate pair data. Using Geneious 8.1 and its Map to Reference feature the cleaned reads were assembled to each of the template scaffolds. BAM files were exported for each paired-end and mate pair assembly. SAMtools [114] was used to merge BAM files to create an assembly with both types of reads. This merged BAM file was imported into Geneious 8.1 where consensus sequences were determined for each scaffold using majority calling to limit the number of ambiguities. GTF files for each scaffold used were retrieved from FlyBase version FB2015_02 [112]. These annotations were transferred to each of the new genomes by aligning each assembled genome scaffold to the reference genome scaffold using Mauve Genome Alignment [115] with default settings except for selecting assume collinear genomes. After alignment, annotations were transferred from the reference to the new assembly. The resulting scaffolds were exported in GenBank format. Using the EMBOSS program, extractfeat [116], CDS sequences were extracted from the assembled scaffolds. Sequence files for each gene were concatenated and then aligned using the default settings of the aligner Muscle 3.8.31 [117]. Only the longest transcript for each gene was used as some genes have multiple splice variants.

### Molecular evolution analysis

To generate substitution counts for filtering, the software KaKs Calculator 1.2 [45] was used. Files of aligned genes were converted to AXT format using the Perl script parseFastaIntoAXT.pl including in the package. After conversion each gene was run through the software using the NG method [118]. The output files for each loci were concatenated and then imported into JMP 10 for filtering.

Values for ω were calculated using codeml part of the PAML 4.9 package [46]. Aligned genes were converted to PHYLIP format using BioPerl [119]. As PAML requires a phylogenetic tree to be provided for its calculations a neighbor joining tree was constructed in MEGA 5 [120]. This was done by concatenating all exons from each population and then aligning them using Mauve Genome Alignment [115]. The alignment was converted to MEG format using MEGA and a neighbor joining tree was built using the default settings. The tree was exported in newick format for use by PAML. Genes were removed from analysis if they were not divisible by three, these genes were manually screened and if alignment errors appeared to be the cause, these were manually corrected. Screening was done for stop codons within the sequences by translating the DNA sequence to protein sequence with Transeq, part of the EMBOSS package [116] and any genes with internal stop codons were removed.

Using the BioPython PAML module [121], control files were built for each gene alignment with default values taken except codon frequency was set to F3×4. Site-class models 0, 7, and 8 were used to calculate the ω values [122–124]. Model 0 is a single ratio based omega value for the entire gene. Model 7 is a null model with 10 classes, which does not allow for positive selection while model 8 adds an additional class that allows for positive selection. Both the ω values and log likelihood values were extracted from each output file and the data was organized in Microsoft Excel. If model 8 significantly better fits the data this is evidence of positive selection [46]. Significance values were found by taking the difference between the log likelihood values of the two outputs and multiplying them by two. This value was then compared a chi-square distribution to find *P* values for each gene. Genes with less than five total substitutions as determined by KaKs Calculator [45] were filtered out and not considered. This was done to help deal with the low power of these methods when there are very few changes between the populations. Genes with few changes are more likely to cause the software to either return an undefined result or to reach the maximum ω the software allows. In addition, genes with either no nonsynonymous or no synonymous changes were also removed. This yielded a total of 9,087 genes that were used in the analysis. Histograms of a log2 transformation of the ω values were produced using JMP 10. A comparison between the log2 transformations of the NG Ka/Ks and the omega value from model 0 of codeml was generated with JMP 10.

The length of each gene’s coding sequence was extracted from the PHYLIP sequence headers. This was to determine if genes with longer length have significantly different omega values. Genes were binned based on length and an ANOVA with post-hoc Tukey test using JMP 10 was used to compare length bins for significance. Intron data was extracted from the reference genome annotation using Geneious 8.1. Based on this, genes were binned based on the number of exons. ANOVA with post-hoc Tukey test in JMP 10 compared the bin sets for significant difference in omega. To determine if there was a significant difference in omega between genes present on each Muller element ANOVA with post-hoc Tukey test was used in JMP 10 to compare omega value distribution on each element.

### Expression analysis

Previous transcriptional studies provided differential expression data for cactus host shifts [43] and between populations [44]. Loci that were found to be significant with codeml model 7 and 8 were removed from this analysis. The model 0 omega for loci with a FDR significance greater than 0.001 for third-instar larva from the *D. mojavensis* Sonora population that were raised on agria cactus rot was compared to non-significant loci using ANOVA in JMP 10. Comparison of model 0 omega between FDR significant loci and non-significant loci was also done for differential expression between third-instar larva of the four host populations with ANOVA in JMP 10.

To explore the relationship between omega and gene expression level RNAseq data from [47] was retrieved for whole male and female *D. mojavensis* flies as aligned BAM files. Differential expression was calculated by using edgeR [125] to look for genes with significantly higher male or female expression. Box plots of omega model 0 for genes with significant male or female expressed genes as well as genes without sex based expression were compared using ANOVA with post-hoc Tukey test in JMP 10. Average adjusted (+0.25) log2 RPKM of non-sex biased genes was plotted against log2 omega model 0 and linear regression was performed on the data with JMP 10.

### Gene ontology terms analysis

Network graphs were generated using Cytoscape 3.2.1 [126] with the add-on app ClueGO 2.2.5 [127]. GO term and KEGG pathway data used was from the June 2016 release. The custom *D. melanogaster* reference set was used for analysis. Both the TOP10 and PAML-FDR genes were run on, biological processes, molecular function and KEGG terms. Data for GO term summary tables was retrieved from FlyBase version FB2017_06 *D. melanogaster* release 6.19 [112]. For each *D. mojavensis* gene with a *D. melanogaster* ortholog, GO term summaries were phrased from the FlyBase GO Summary Ribbons for molecular function and biological process. Clustering done with JMP 10 using the Ward method and 15 groups allowed.

## Supporting information

Functional clustering Table S11

## Abbreviations

2L: Left arm of 2nd chromosome in *D. melanogaster*
2R: Right arm of 2nd chromosome in *D. melanogaster*
3L: Left arm of 3rd chromosome in *D. melanogaster*
3R: Right arm of 3rd chromosome in *D. melanogaster*
ACP: Accessory gland protein
ANOVA: Analysis of Variance
BAM: Binary Alignment Map
CDS: Coding sequence
EMBOSS: European Molecular Biology Open Software Suite
FDR: False Discovery Rate
GO: Gene Ontology
Gst: Glutathione S-transferase
Ka: number of nonsynonymous substitution per nonsynonymous site
kb: Kilobase
KEGG: Kyoto Encyclopedia of Genes and Genomes
Ks: number of synonymous substitution per synonymous site
MEGA: Molecular Evolutionary Genetics Analysis software
PAML: Phylogenetic Analysis of Maximum Likelihood program
PAML-FDR: PAML significant loci post-FDR correction
PHYLIP: Phylogeny Inference Package
RPKM: Reads Per Kilobase per Million mapped reads
TOP10: Loci with ω values in the top 10% of the distribution

## Availability of data and materials

The datasets supporting the conclusions of this article are available in the NCBI Sequence Read Archive (SRA) under the accession number PRJNA530196 (http://XXX) and OSF (https://doi.org/XXXXX). Additionally, datasets supporting the conclusions of this article are included within the article its additional files.

## Competing interests

The authors declare that they have no competing interests.

## Authors’ contributions

CWA performed the assembly and analysis of the genomic data and was involved in the writing of the manuscript. LMM conceived of and designed the study, was involved in the analysis and the writing of the manuscript. All authors read and approved the final manuscript.

## Acknowledgements

The authors greatly acknowledge the work of Laurel Brandsmeier in this project. This work was supported by a Junior Faculty Distinguished Research award from the University of Alabama in Huntsville and partly supported by a grant from the National Science Foundation (DEB-1219387 and IOS-1557697 to LMM).

