## Supplemental Tables and Figures for "Genomic analysis of the four ecologically distinct cactus host populations of *Drosophila mojavensis*"

**Table S2** Pairwise Tukey HSD of $\log_2 \omega$ between Muller elements

| Element | Element | Difference | Std Err Dif | Lower CL | Upper CL | *P* Value |
|---|---|---|---|---|---|---|
| F | C | 1.078745 | 0.1890123 | 0.539991 | 1.617498 | <.0001 |
| F | D | 1.033424 | 0.1892502 | 0.493993 | 1.572856 | <.0001 |
| F | A | 1.014120 | 0.1906562 | 0.470681 | 1.557560 | <.0001 |
| F | B | 0.714138 | 0.1892686 | 0.174654 | 1.253622 | 0.0022 |
| F | E | 0.667516 | 0.1883892 | 0.130538 | 1.204493 | 0.0053 |
| E | C | 0.411229 | 0.0480779 | 0.274189 | 0.548268 | <.0001 |
| E | D | 0.365908 | 0.0490047 | 0.226227 | 0.505590 | <.0001 |
| B | C | 0.364607 | 0.0514158 | 0.218053 | 0.511160 | <.0001 |
| E | A | 0.346605 | 0.0541814 | 0.192168 | 0.501041 | <.0001 |
| B | D | 0.319287 | 0.0522834 | 0.170260 | 0.468313 | <.0001 |
| B | A | 0.299983 | 0.0571640 | 0.137045 | 0.462921 | <.0001 |
| A | C | 0.064624 | 0.0563096 | -0.095878 | 0.225127 | 0.8613 |
| E | B | 0.046622 | 0.0490758 | -0.093262 | 0.186506 | 0.9333 |
| D | C | 0.045320 | 0.0513479 | -0.101040 | 0.191681 | 0.9508 |
| A | D | 0.019304 | 0.0571029 | -0.143460 | 0.182068 | 0.9994 |

**Table S3** Pairwise Tukey HSD of $\log_2 \omega$ between different coding gene length bins.

| Length Bin | Length Bin | Difference | Std Err Dif | Lower CL | Upper CL | *P* Value |
|---|---|---|---|---|---|---|
| 1 | 4 | 1.187473 | 0.0668196 | 1.00516 | 1.369784 | <.0001 |
| 1 | 5 | 1.147404 | 0.0627699 | 0.97614 | 1.318665 | <.0001 |
| 1 | 3 | 1.051908 | 0.0494634 | 0.91695 | 1.186864 | <.0001 |
| 1 | 2 | 0.767654 | 0.0384280 | 0.66281 | 0.872501 | <.0001 |
| 2 | 4 | 0.419819 | 0.0648992 | 0.24275 | 0.596890 | <.0001 |
| 2 | 5 | 0.379750 | 0.0607215 | 0.21408 | 0.545422 | <.0001 |
| 2 | 3 | 0.284254 | 0.0468366 | 0.15647 | 0.412043 | <.0001 |
| 3 | 4 | 0.135565 | 0.0719849 | -0.06084 | 0.331968 | 0.3264 |
| 3 | 5 | 0.095496 | 0.0682424 | -0.09070 | 0.281688 | 0.6281 |
| 5 | 4 | 0.040069 | 0.0817022 | -0.18285 | 0.262985 | 0.9883 |

Bin 1 = < 1 Kb, Bin 2 = 1-2 Kb, Bin 3 = 2-3 Kb, Bin 4 = 3-4 Kb, Bin 5 >4 Kb.

**Table S4** Pairwise Tukey HSD of $\log_2 \omega$ between different exon number bins.

| Exon Bin | Exon Bin | Difference | Std Err Dif | Lower CL | Upper CL | *P* Value |
|---|---|---|---|---|---|---|
| 2 | 7 | 0.9818659 | 0.0507210 | 0.832289 | 1.131443 | <.0001 |
| 1 | 7 | 0.7994915 | 0.0544689 | 0.638862 | 0.960121 | <.0001 |
| 3 | 7 | 0.7401690 | 0.0529489 | 0.584022 | 0.896316 | <.0001 |
| 2 | 5 | 0.6570861 | 0.0652903 | 0.464544 | 0.849628 | <.0001 |
| 2 | 6 | 0.6087085 | 0.0717851 | 0.397013 | 0.820404 | <.0001 |
| 2 | 4 | 0.5297340 | 0.0572028 | 0.361042 | 0.698426 | <.0001 |
| 1 | 5 | 0.4747116 | 0.0682426 | 0.273463 | 0.675960 | <.0001 |
| 4 | 7 | 0.4521319 | 0.0569040 | 0.284321 | 0.619943 | <.0001 |
| 1 | 6 | 0.4263340 | 0.0744805 | 0.206690 | 0.645978 | <.0001 |
| 3 | 5 | 0.4153892 | 0.0670357 | 0.217700 | 0.613079 | <.0001 |
| 6 | 7 | 0.3731574 | 0.0715473 | 0.162163 | 0.584151 | <.0001 |
| 3 | 6 | 0.3670116 | 0.0733762 | 0.150624 | 0.583399 | <.0001 |
| 1 | 4 | 0.3473595 | 0.0605508 | 0.168794 | 0.525925 | <.0001 |
| 5 | 7 | 0.3247798 | 0.0650287 | 0.133009 | 0.516550 | <.0001 |
| 3 | 4 | 0.2880371 | 0.0591872 | 0.113493 | 0.462581 | <.0001 |
| 2 | 3 | 0.2416969 | 0.0532698 | 0.084603 | 0.398791 | 0.0001 |
| 2 | 1 | 0.1823745 | 0.0547809 | 0.020825 | 0.343924 | 0.0153 |
| 4 | 5 | 0.1273521 | 0.0702016 | -0.079674 | 0.334378 | 0.5384 |
| 4 | 6 | 0.0789745 | 0.0762795 | -0.145975 | 0.303924 | 0.9459 |
| 1 | 3 | 0.0593224 | 0.0568499 | -0.108329 | 0.226974 | 0.9439 |
| 6 | 5 | 0.0483776 | 0.0825179 | -0.194969 | 0.291724 | 0.9972 |

Bin 1 = 1 exon, Bin 2 = 2 exons, Bin 3 = 3 exons, Bin 4 = 4 exons, Bin 5 = 5 exons, Bin 6 = 6 exons, Bin 7 > 7 exons.

**Table S5** Pairwise Tukey HSD of $\log_2 \omega$ between different expression categories.

| Category | Category | Difference | Std Err Dif | Lower CL | Upper CL | *P* Value |
|---|---|---|---|---|---|---|
| Male bias | Female bias | 0.5461726 | 0.0494397 | 0.4302817 | 0.6620635 | <.0001 |
| Male bias | Unbiased | 0.3344762 | 0.0397754 | 0.2412392 | 0.4277131 | <.0001 |
| Unbiased | Female bias | 0.2116964 | 0.0425320 | 0.1119977 | 0.3113952 | <.0001 |

**Table S6** Pairwise Tukey HSD of $\log_2 \omega$ between different expression categories.

| Category | Category | Difference | Std Err Dif | Lower CL | Upper CL | *P* Value |
|---|---|---|---|---|---|---|
| 1 | 2 | 0.5368926 | 0.0386429 | 0.446310 | 0.6274750 | <.0001 |
| 1 | 3 | 0.4194892 | 0.0698346 | 0.255791 | 0.5831875 | <.0001 |
| 3 | 2 | 0.1174035 | 0.0643864 | -0.033524 | 0.2683308 | 0.1620 |

RPKM Category 1 = <-2,2.4>, RPKM Category 2 = <2.4,6.8>, RPKM Category 3 = <6.8,11.2>

**Table S7** ANOVA results of $\log_2 \omega$ differences between loci differentially expressed as a result of cactus rearing and loci differentially expressed across populations.

| Source | DF | Sum of Squares | Mean Square | F Ratio | Prob > F |
|---|---|---|---|---|---|
| Cactus Effect | 1 | 135.11 | 135.11 | 60.64 | <0.0001 |
| Error | 8317 | 18531.41 | 2.23 | | |
| Total | 8318 | 18666.52 | | | |
| Population Effect | 1 | 116.64 | 116.64 | 52.31 | <0.0001 |
| Error | 8319 | 18549.44 | 2.23 | | |
| Total | 8320 | 18666.08 | | | |

PAML significant loci removed. Cactus effect analysis composed of 8319 loci, population effect analysis composed of 8321 loci.

**Table S8** List of overrepresented GO terms among databases

| Gene dataset | Term dataset | Term | *P* value [a] |
|---|---|---|---|
| TOP10 | Molecular process | Serine-type peptidase activity | 0.0000011 |
| | | Endopeptidase activity | 0.012 |
| | | Serine-type endopeptidase activity | 0.0000011 |
| | | tRNA-specific ribonuclease activity | 0.0074 |
| TOP10 | KEGG | Oxidative phosphorylation | 0.01 |
| PAML-FDR | Biological function | DNA metabolic process | |

[a] *P* values are FDR corrected

**Table S9** Karyotype of the three *D. mojavensis* lines sequenced and of the template genome line.

| | Muller E (2$^{nd}$ chromosome) | Muller B (3$^{rd}$ chromosome) |
|---|---|---|
| Catalina Island (15081-1352.00)[a] | st/st | $f^2/f^2$ |
| Mojave (MJANZA 402-8)[b] | st/st | st/st |
| Baja California (MJBC 155)[b] | $q^5/q^5$ | st/st |
| Sonora (MJ 122)[b] | $q^5/q^5$ | $f^2/f^2$ |

[a] Karyotype data from [1]. [b] Data from visual inspection of polytene chromosome squashes and *de novo* assembly of genomes (unpublished data, Matzkin).

**Table S10** List of reference genome scaffolds used for template assembly.

| *D. mojavensis* chromosome | Scaffold ID | Homologous *D. melanogaster* arm | | Size (bp) |
|---|---|---|---|---|
| Muller A Chromosome 1 | 1786 | X | | 4,599 |
| | 6308 | X | | 3,356,042 |
| | 6314 | X | | 47,931 |
| | 6328 | X | | 4,453,435 |
| | 6359 | X | | 4,525,533 |
| | 6473 | X | | 16,943,266 |
| | 6482 | X | | 2,735,782 |
| | 6541 | X | | 2,543,558 |
| | 6657 | X | | 3,686 |
| | 6660 | X | | 7,398 |
| | | | Total Size | 34,621,230 |
| Muller B Chromosome 3 | 587 | 2L | | 1,541 |
| | 6005 | 2L | | 1,209 |
| | 6200 | 2L | | 24,743 |
| | 6500 | 2L | | 32,352,404 |
| | 6583 | 2L | | 2,108 |
| | 6584 | 2L | | 818 |
| | 6587 | 2L | | 1,717 |
| | 6615 | 2L | | 6,841 |
| | 6646 | 2L | | 1,733 |
| | | | Total Size | 32,393,114 |
| Muller C Chromosome 5 | 1740 | 2R | | 7,530 |
| | 3877 | 2R | | 1,968 |
| | 6071 | 2R | | 80,338 |
| | 6496 | 2R | | 26,866,924 |
| | | | Total Size | 26,956,760 |
| Muller D Chromosome 4 | 4729 | 3L | | 8,237 |
| | 4734 | 3L | | 13,141 |
| | 4739 | 3L | | 8,224 |
| | 5947 | 3L | | 5,469 |
| | 5978 | 3L | | 5,259 |
| | 6654 | 3L | | 2,564,135 |
| | 6680 | 3L | | 24,764,193 |
| | | | Total Size | 27,368,658 |
| Muller E Chromosome 2 | 1710 | 3R | | 6,556 |
| | 1718 | 3R | | 5,511 |
| | 4072 | 3R | | 13,853 |
| | 4889 | 3R | | 3,248 |
| | 6111 | 3R | | 58,398 |
| | 6368 | 3R | | 49,292 |
| | 6540 | 3R | | 34,148,556 |
| | | | Total Size | 34,285,414 |
| Muller F Chromosome 6 | 6498 | 4 | | 3,408,170 |
| | | | Total Size | 3,408,170 |
| Unmapped | 6499 | | | 421,677 |

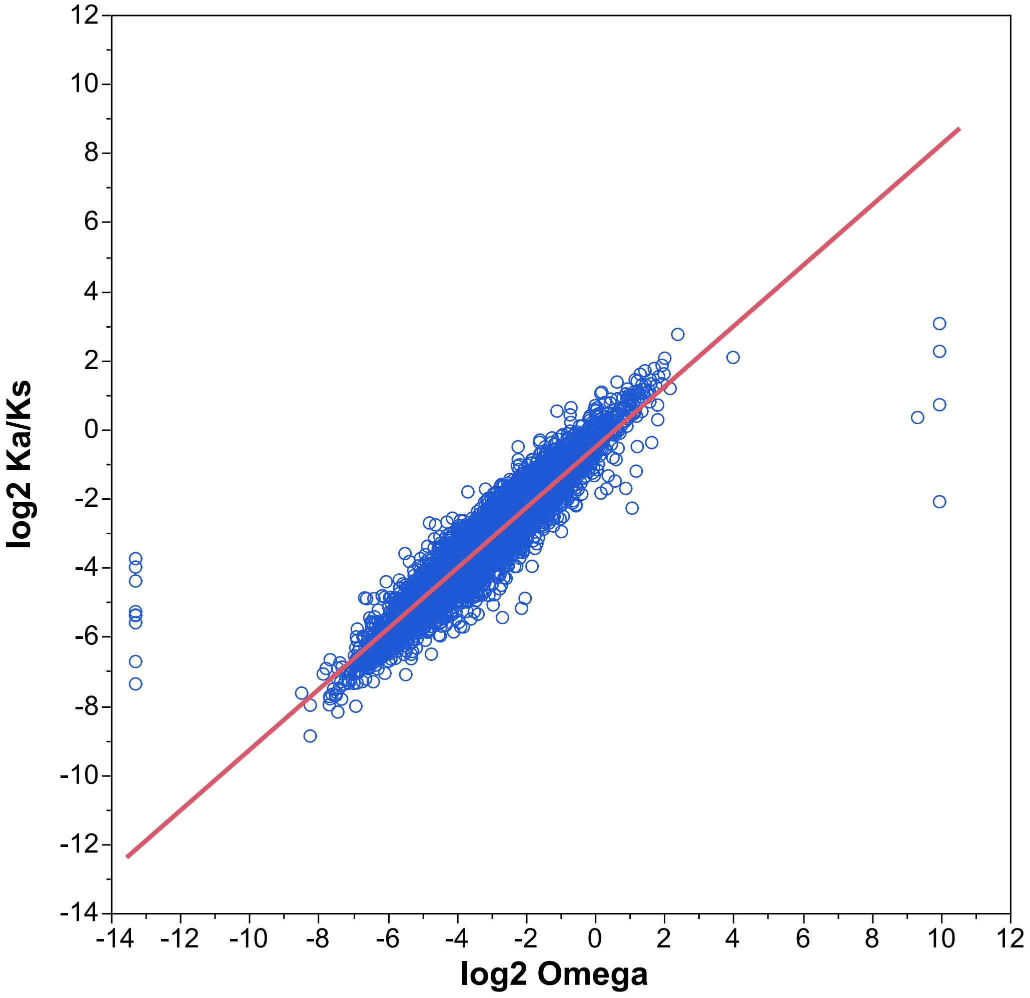

**Fig. S1** Relationship between $\log_2$ Ka/Ks values estimated via the KaKs Calculator and $\log_2$ ω values estimated via Codeml among the 9,087 post filter genes ($r^2 = 0.88$, $P < 0.001$).

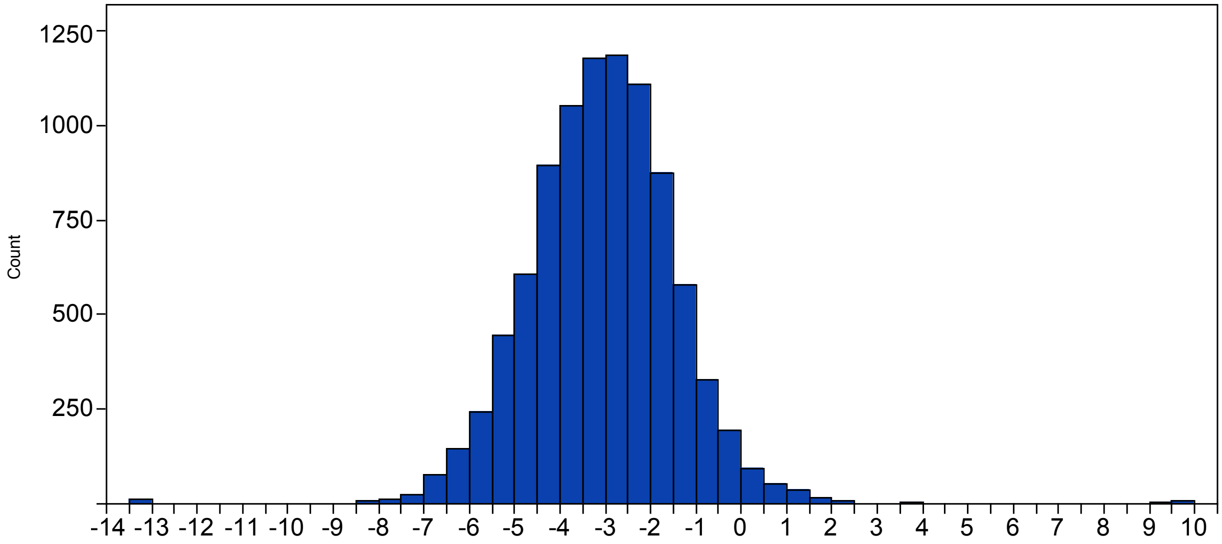

**Fig. S2** Distribution of $\log_2 \omega$ values for all loci that passed filtering (>5 total substitutions, > 0 nonsynonymous substitutions, > 0 synonymous substitutions), a total of 9,087 genes.

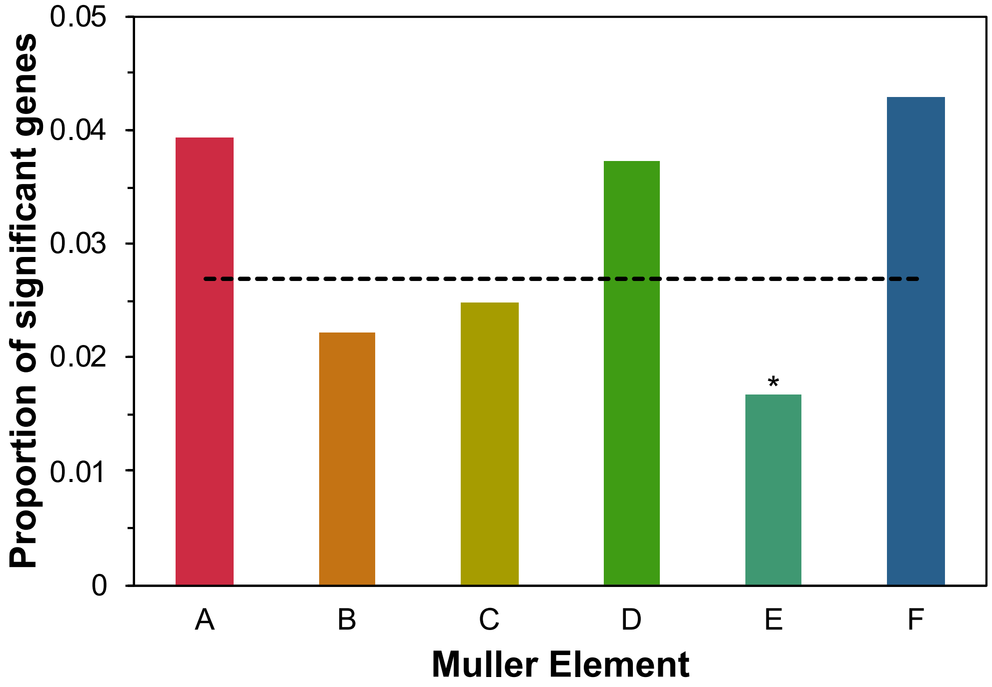

**Fig. S3** Proportion of positive selected (via PAML with FDR correction) loci located in each of the *D. mojavensis* Muller elements. Dashed line indicates the genome wide proportion of positive selected loci (0.0269). Asterisk indicate significance via Fisher's Exact test (* P < 0.05, ** P < 0.01, *** P < 0.001).

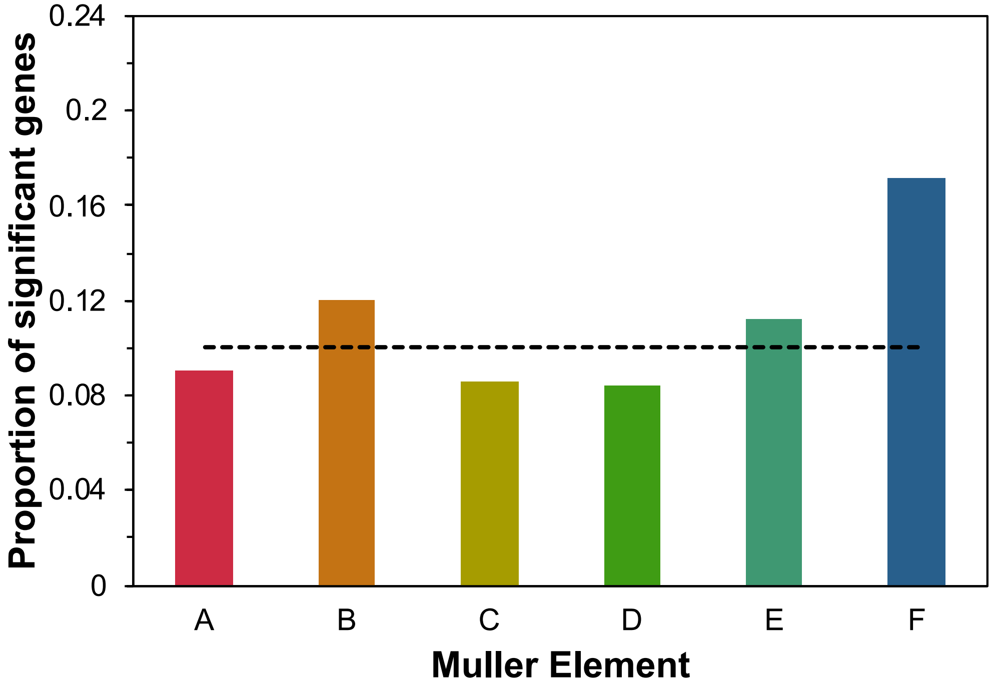

**Fig. S4** Proportion of TOP10 loci located in each of the *D. mojavensis* Muller elements. Dashed line indicates the genome wide proportion of TOP10 loci (0.10). Asterisk indicate significance via Fisher's Exact test (* P < 0.05, ** P < 0.01, *** P < 0.001).

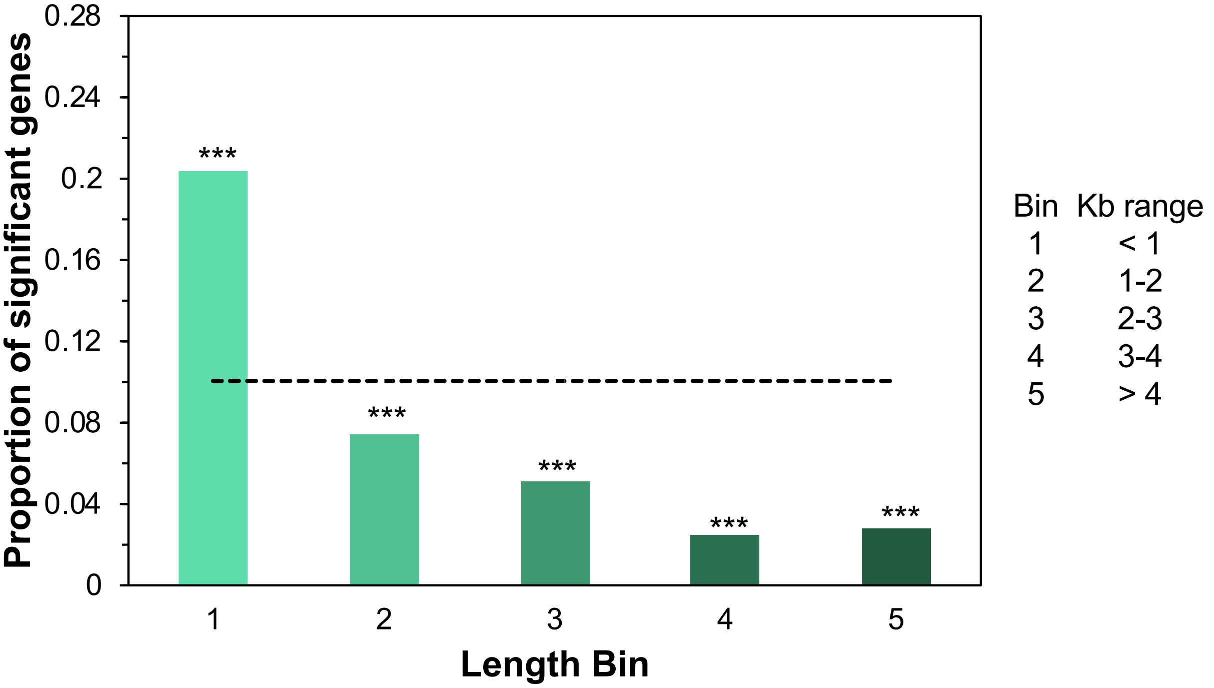

**Fig. S5** Proportion of TOP10 loci in each of five coding length bins. Dashed line indicates the genome wide proportion of TOP10 loci (0.10). Asterisk indicate significance via Fisher's Exact test (* P < 0.05, ** P < 0.01, *** P < 0.001).

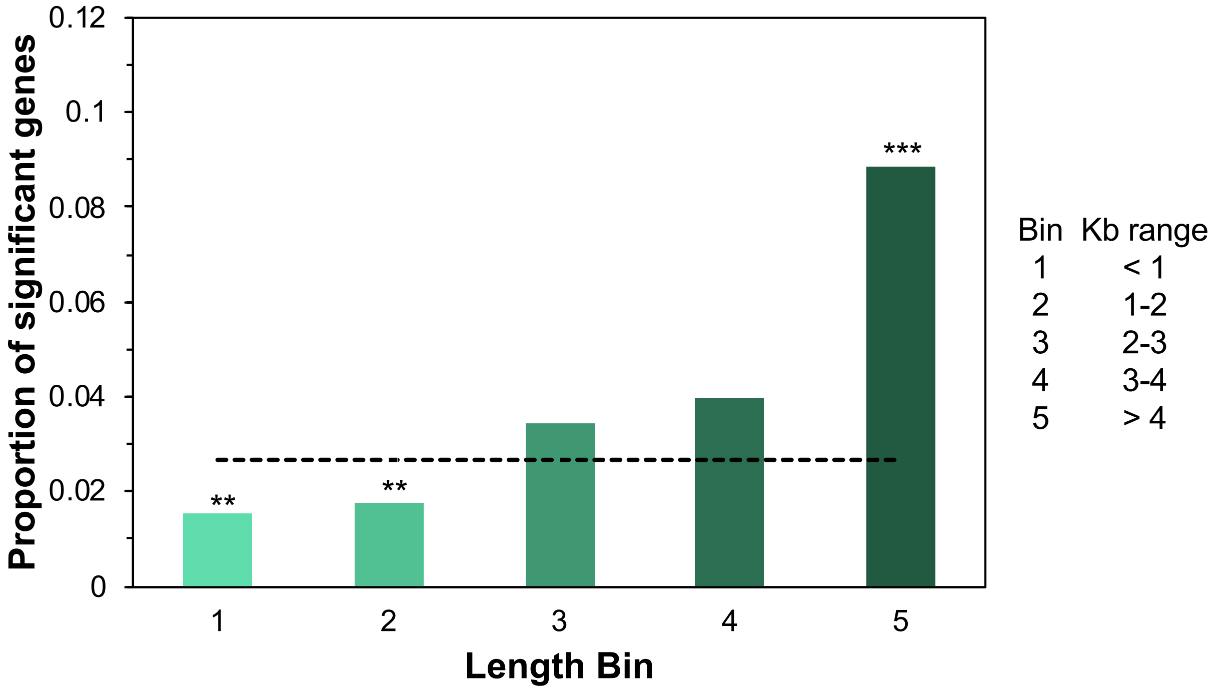

**Fig. S6** Proportion of positive selected (via PAML with FDR correction) loci in each of five coding length bins. Dashed line indicates the genome wide proportion of positive selected loci (0.0269). Asterisk indicate significance via Fisher's Exact test (* P < 0.05, ** P < 0.01, *** P < 0.001).

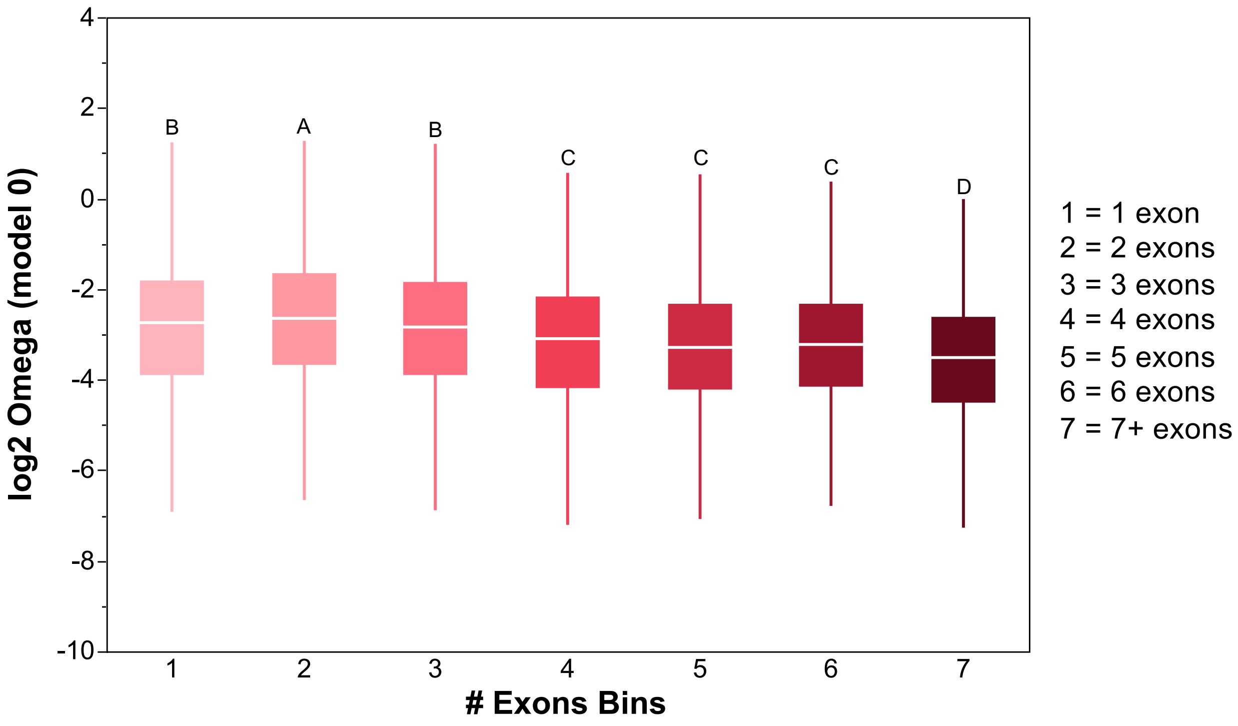

**Fig. S7** Boxplot of $\log_2 \omega$ values for loci in seven different exon number bins. Bins with different letters are significantly different using a Tukey HSD test (see Table S4).

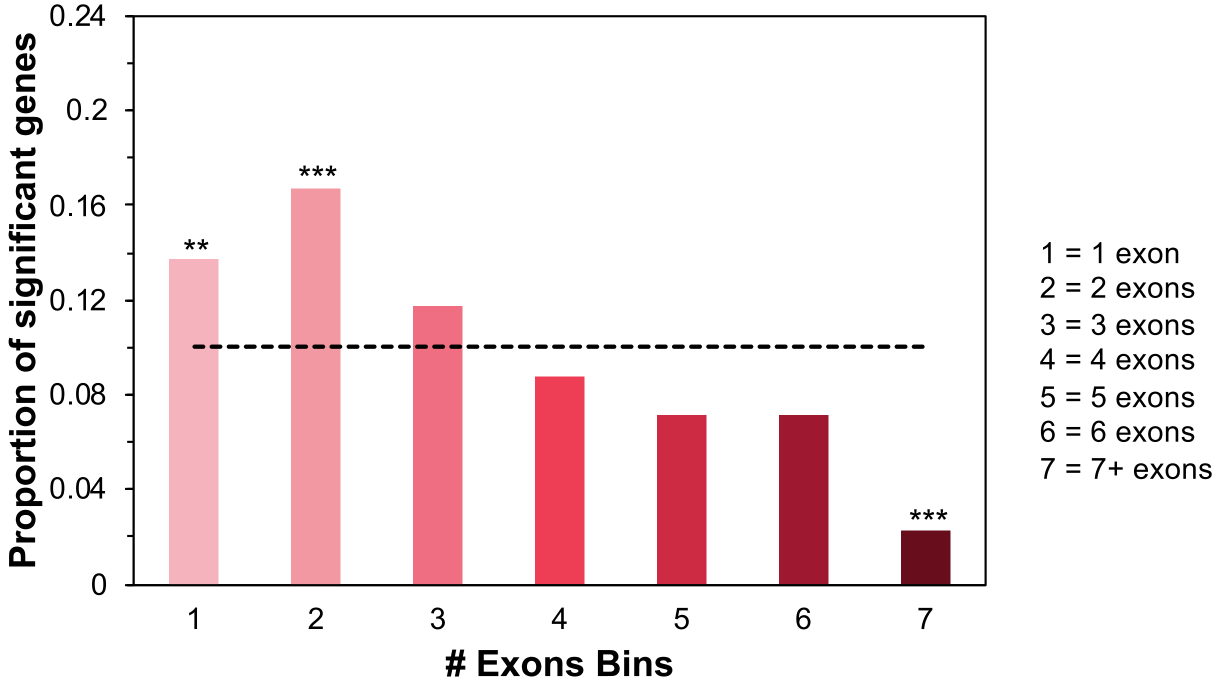

**Fig. S8** Proportion of TOP10 loci in each of seven exon number bins. Dashed line indicates the genome wide proportion of TOP10 loci (0.10). Asterisk indicate significance via Fisher's Exact test (* P < 0.05, ** P < 0.01, *** P < 0.001).

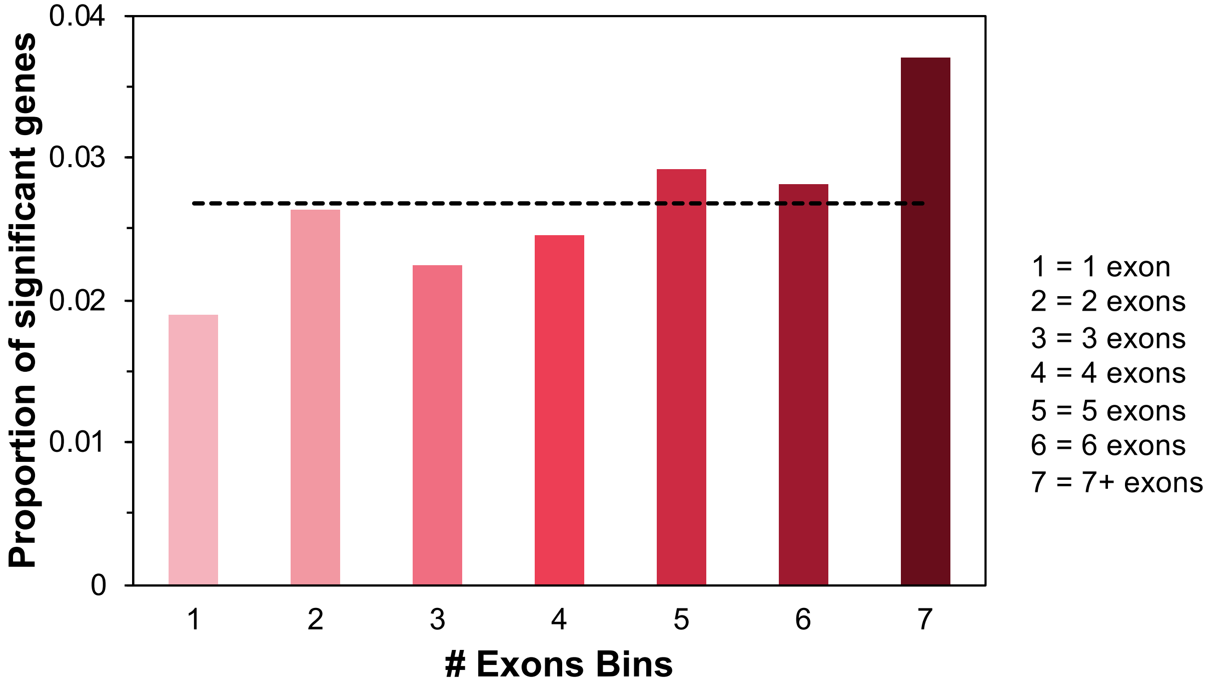

**Fig. S9** Proportion of positive selected (via PAML with FDR correction) loci in each of seven exon number bins. Dashed line indicates the genome wide proportion of positive selected loci (0.0269). Asterisk indicate significance via Fisher's Exact test (* P < 0.05, ** P < 0.01, *** P < 0.001).

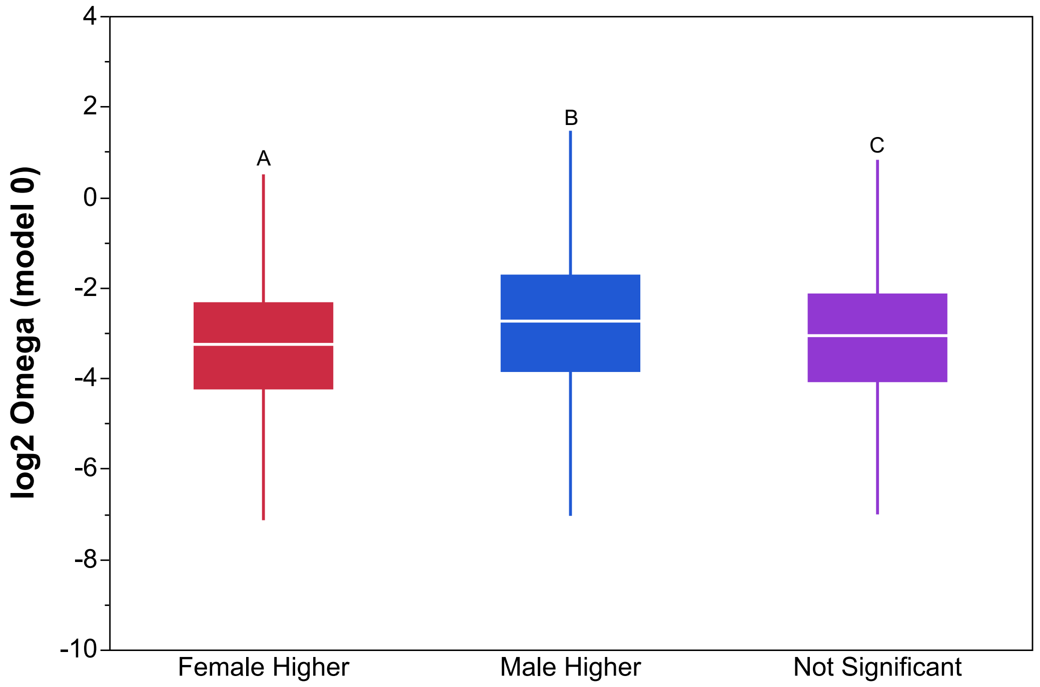

**Fig. S10** Boxplot of $\log_2$ ω values for loci that show female-bias, male-bias or unbiased gene expression. Classes with different letters are significantly different using a Tukey HSD test (see Table S5). Gene expression data is from [2].

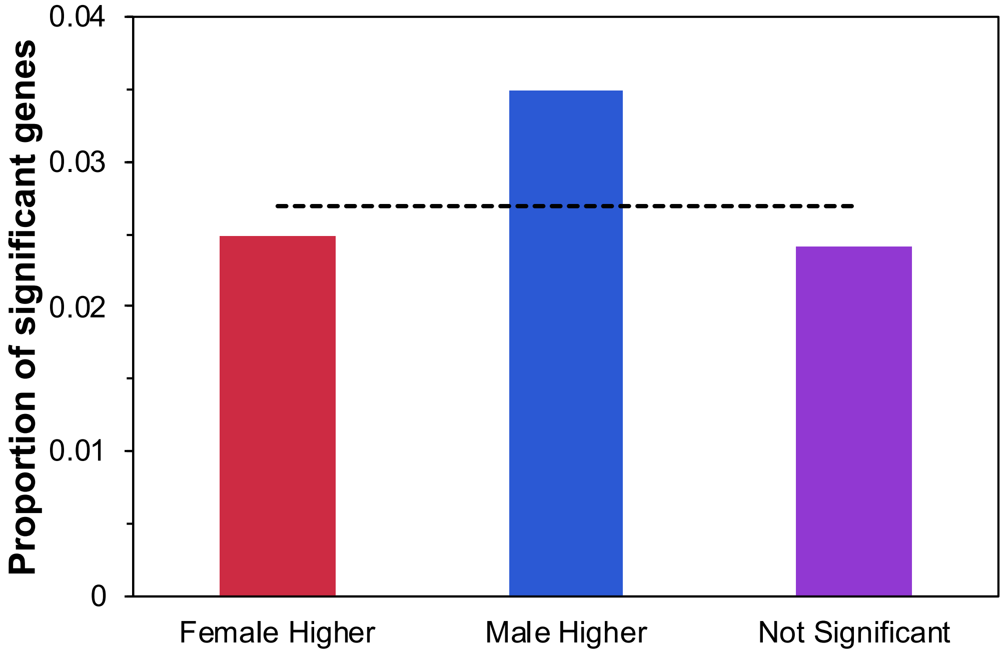

**Fig. S11** Proportion of positive selected (via PAML with FDR correction) loci that show female-bias, male-bias or unbiased gene expression. Dashed line indicates the genome wide proportion of positive selected loci (0.0269). Gene expression data is from [2]. Asterisk indicate significance via Fisher's Exact test (* P < 0.05, ** P < 0.01, *** P < 0.001).

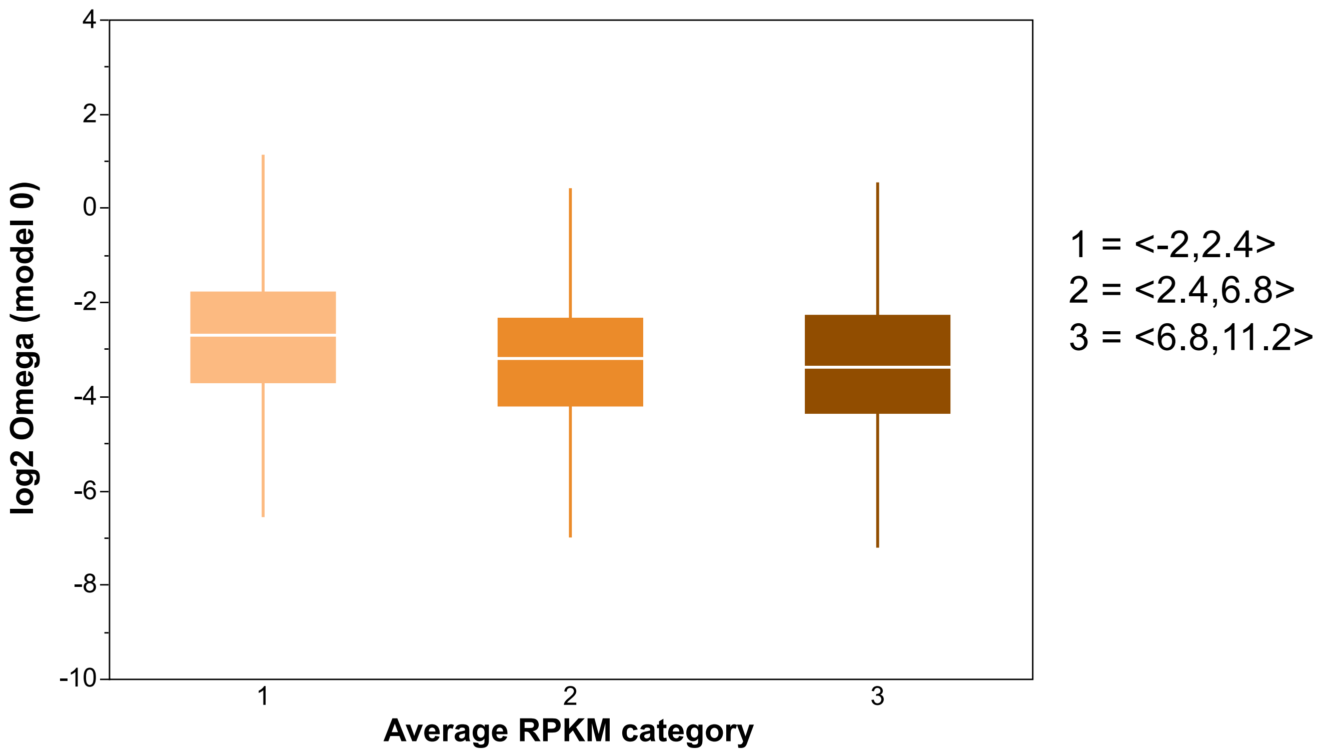

**Fig. S12** Boxplot of $log_2$ ω values for loci in three different expression level categories. Categories with different letters are significantly different using a Tukey HSD test (see Table S6). Gene expression data is from [2]. Female and male biased genes were removed, only 5,101 genes.

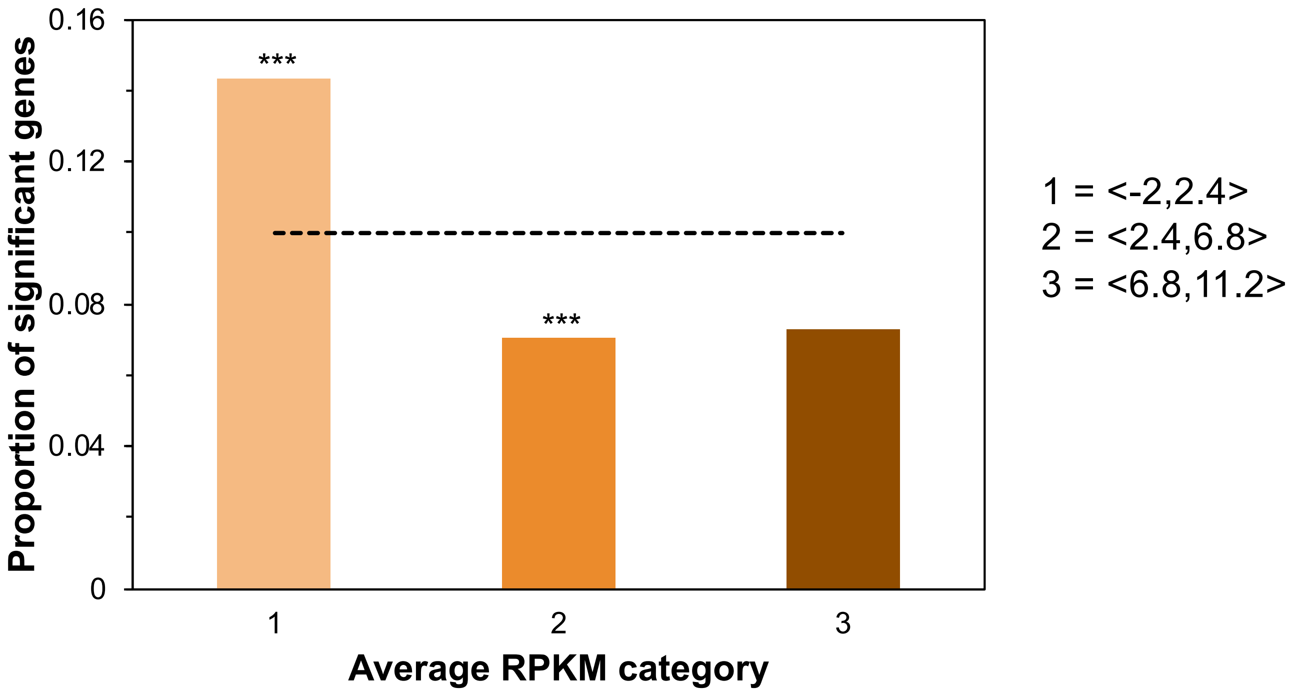

**Fig. S13** Proportion of TOP10 loci in three different expression level categories. Dashed line indicates the genome wide proportion of TOP10 loci (0.10). Gene expression data is from [2]. Female and male biased genes were removed, only 5,101 genes. Asterisk indicate significance via Fisher's Exact test (* P < 0.05, ** P < 0.01, *** P < 0.001).

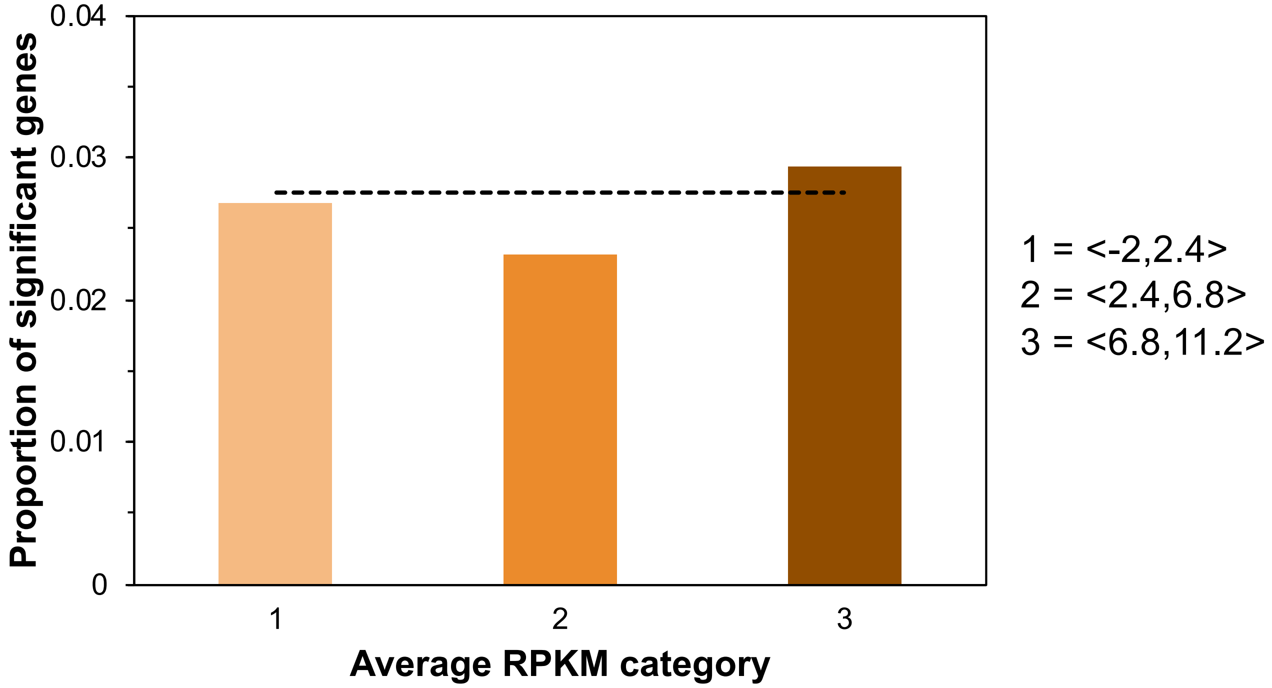

**Fig. S14** Proportion of positive selected (via PAML with FDR correction) loci in three different expression level categories. Dashed line indicates the genome wide proportion of positive selected loci (0.0269). Gene expression data is from [2]. Female and male biased genes were removed, only 5,101 genes. Asterisk indicate significance via Fisher's Exact test (* P < 0.05, ** P < 0.01, *** P < 0.001).

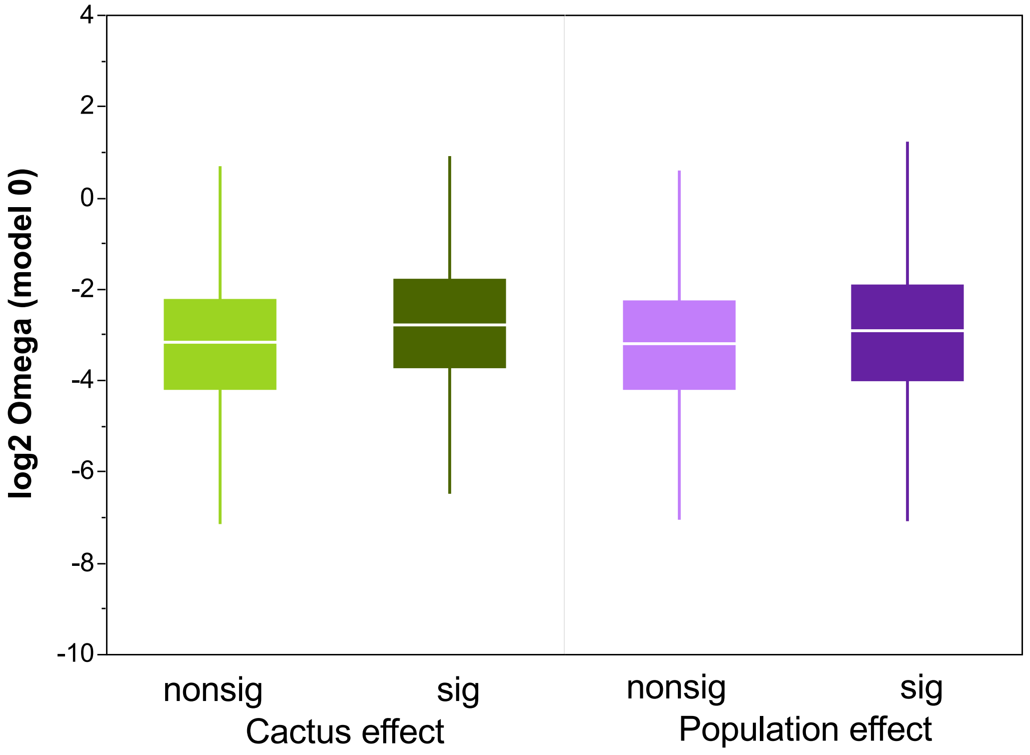

**Fig. S15** Boxplot of $\log_2 \omega$ values for loci differentially expressed as a result of cactus rearing (left) and loci differentially expressed across populations (right). For both panels differences in $\log_2 \omega$ values between differentially and not differentially expressed loci were significant (ANOVA, $P < 0.001$ see Table S7). For both datasets [3, 4] only loci that had a $P < 0.001$ post FDR correction were labeled as differentially expressed. For the integrated analysis PAML significant loci were removed. Cactus effect analysis composed of 8,319 loci, population effect analysis composed of 8,321 loci.

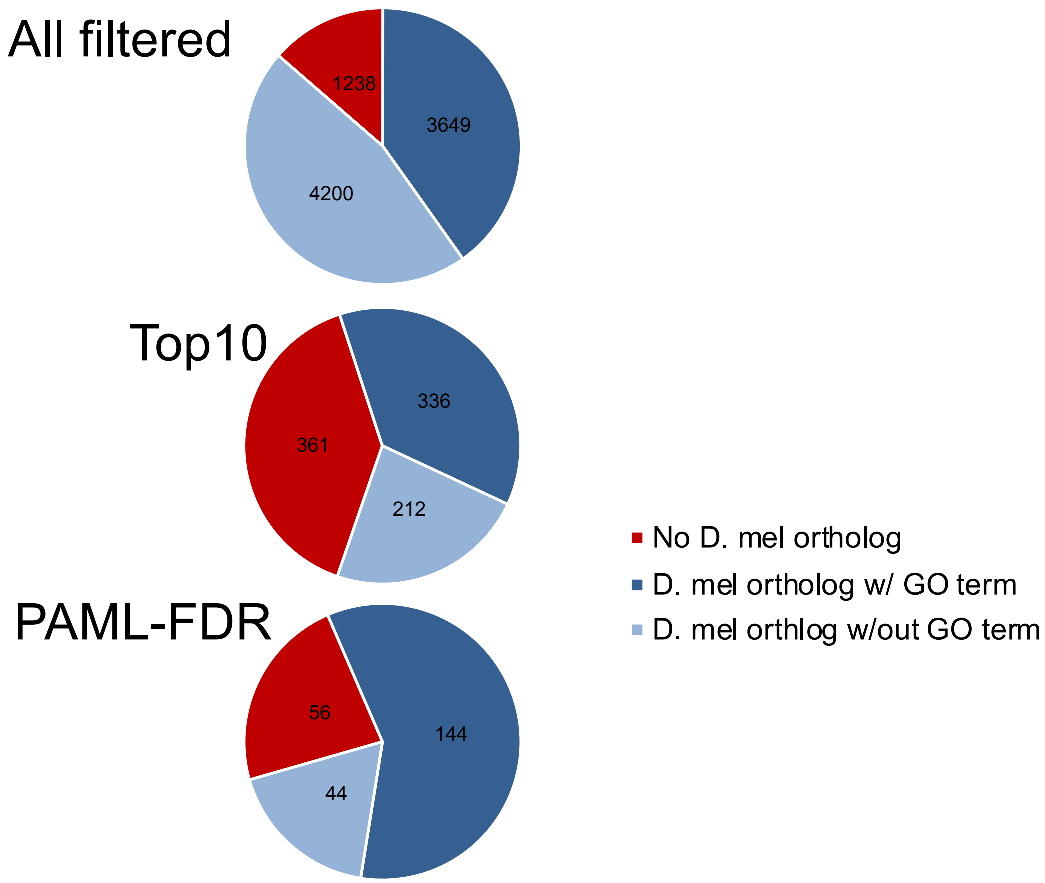

**Fig. S16** Number of orthologous gene calls to the *D. melanogaster* reference genome for the filtered (9,087 genes), TOP10 (909 genes) and PAML-FDR (244 genes).

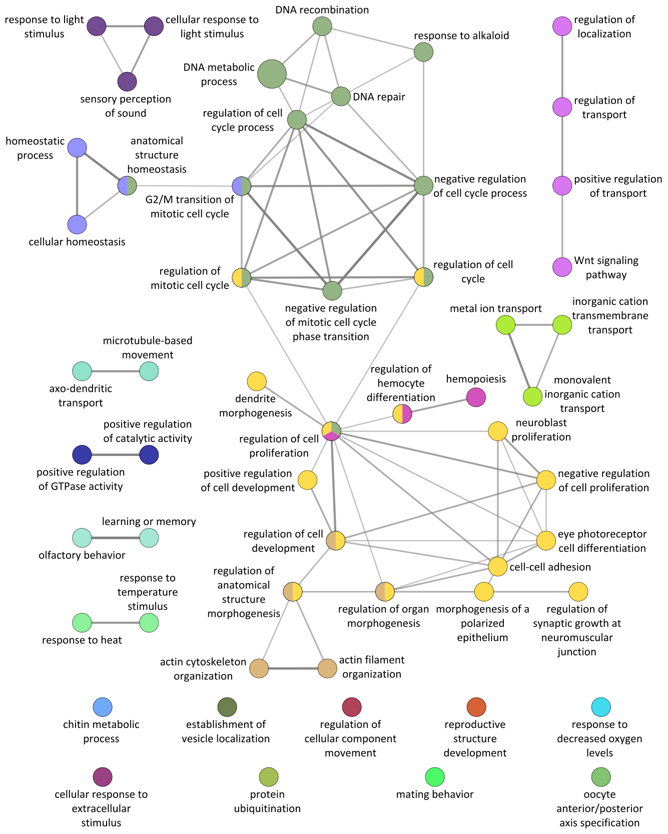

**Fig. S17** Network clustering of Biological Process terms of the positive selected (via PAML with FDR correction) loci. Network clustering was performed using ClueGo using the following parameters: Min GO Level = 3, Max GO Level = 8, All GO Levels = false, Number of Genes = 10, Get All Genes = false, Min Percentage = 4.0, Get All Percentage = false, GO Fusion = true, GO Group = true, Kappa Score Threshold = 0.4 ,Over View Term = SmallestPValue, Group By Kappa Statistics = true, Initial Group Size = 1, Sharing Group Percentage = 50.0

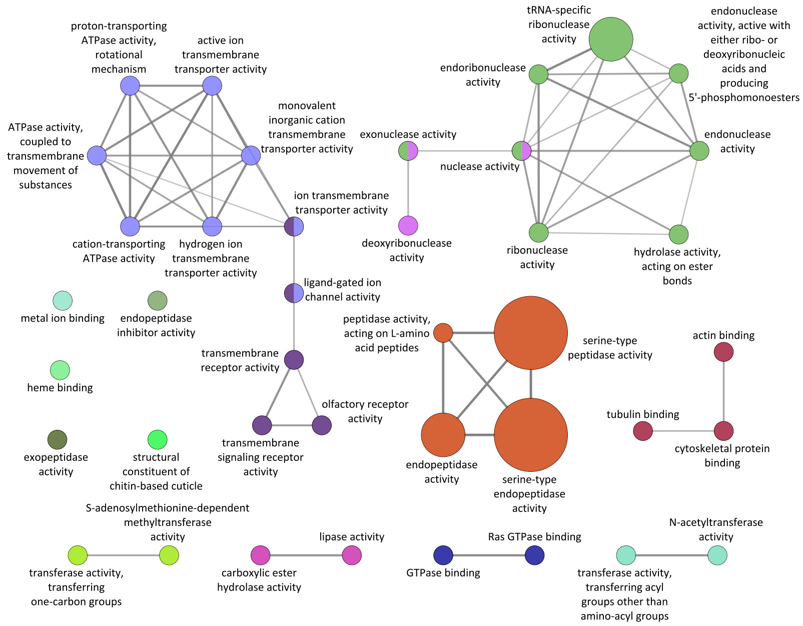

**Fig. S18** Network clustering of Molecular Function terms of the TOP10 loci. Network clustering was performed using ClueGo using the following parameters: Min GO Level = 3, Max GO Level = 8, All GO Levels = false, Number of Genes = 3, Get All Genes = false, Min Percentage = 4.0, Get All Percentage = false, GO Fusion = true, GO Group = true, Kappa Score Threshold = 0.4, Over View Term = SmallestPValue, Group By Kappa Statistics = true, Initial Group Size = 1, Sharing Group Percentage = 50.0

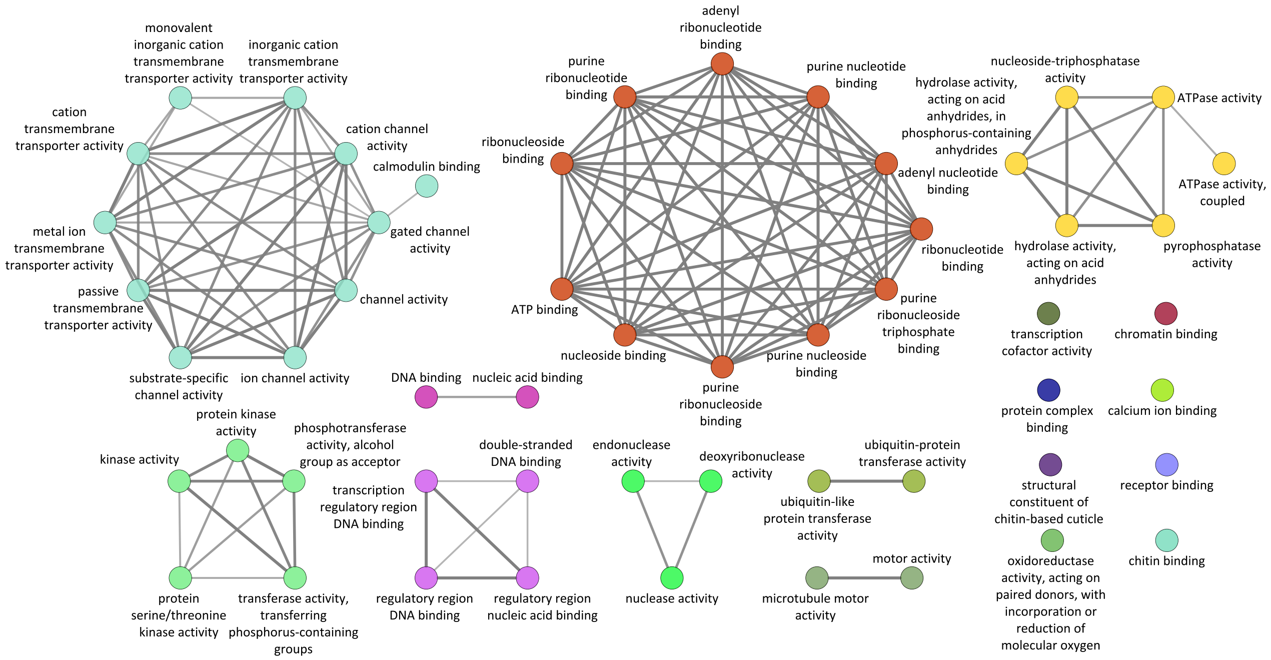

**Fig. S19** Network clustering of Molecular Function terms of the positive selected (via PAML with FDR correction) loci. Network clustering was performed using ClueGo using the following parameters: Min GO Level = 3, Max GO Level = 5, All GO Levels = false, Number of Genes = 10, Get All Genes = false, Min Percentage = 8.0, Get All Percentage = false, GO Fusion = false, GO Group = true, Kappa Score Threshold = 0.4, Over View Term = SmallestPValue, Group By Kappa Statistics = true, Initial Group Size = 1, Sharing Group Percentage = 50.0

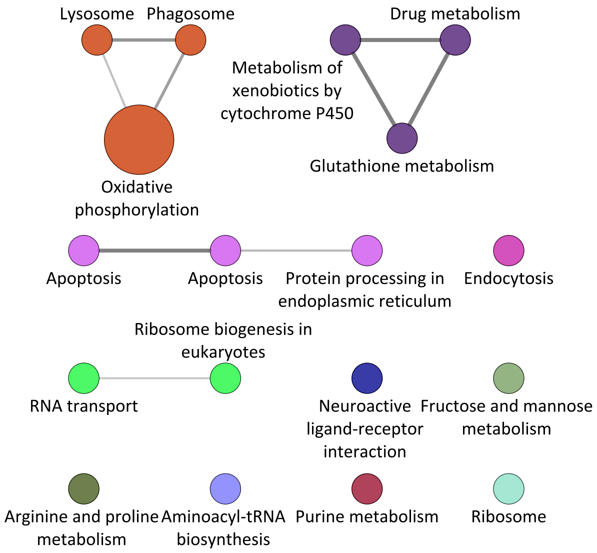

**Fig. S20** Network clustering of KEGG terms of the TOP10 loci. Network clustering was performed using ClueGo using the following parameters: Min GO Level = 3, Max GO Level = 8, All GO Levels = false, Number of Genes = 2, Get All Genes = false, Min Percentage = 1.0, Get All Percentage = false, GO Fusion = true, GO Group = true, Kappa Score Threshold = 0.25, Over View Term = SmallestPValue, Group By Kappa Statistics = true, Initial Group Size = 1, Sharing Group Percentage = 50.0

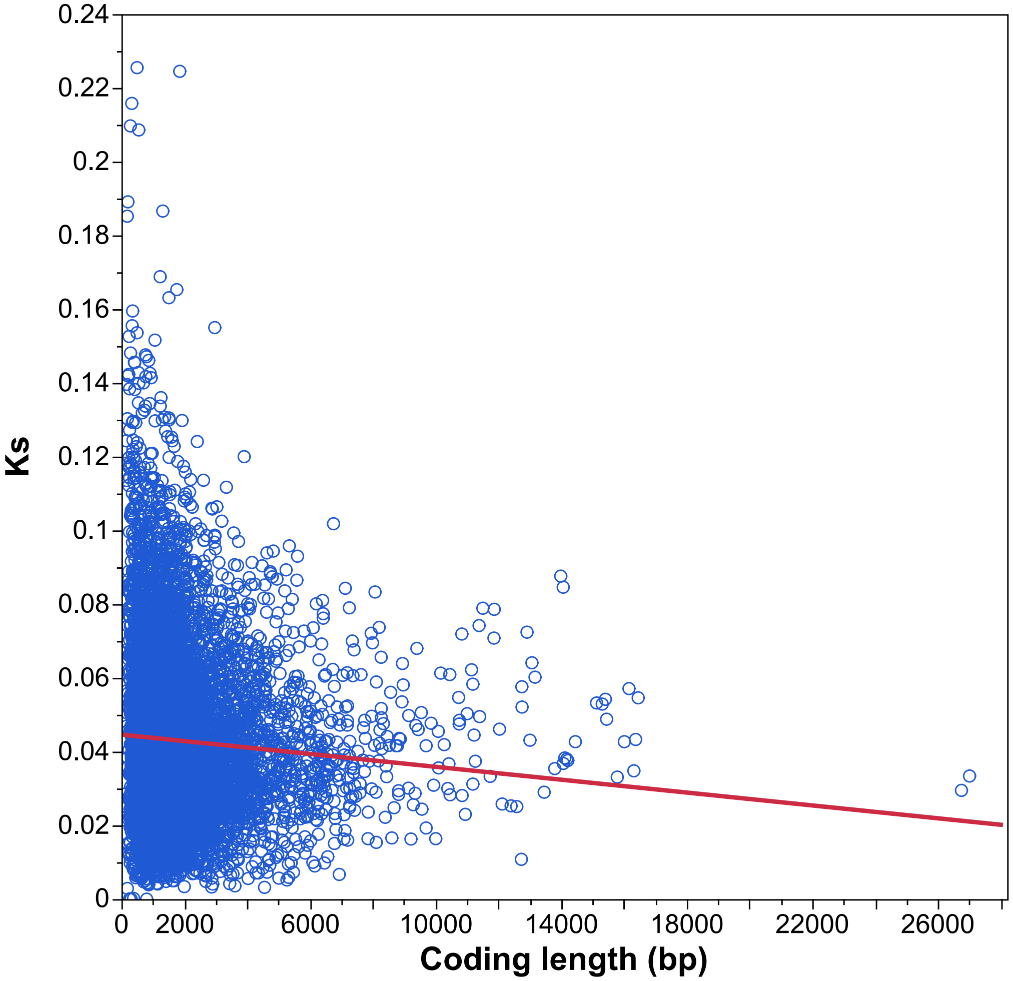

**Fig. S21** Relationship between Ks and gene coding length (9,087 loci) ($r^2 = 0.004$, $P < 0.001$).

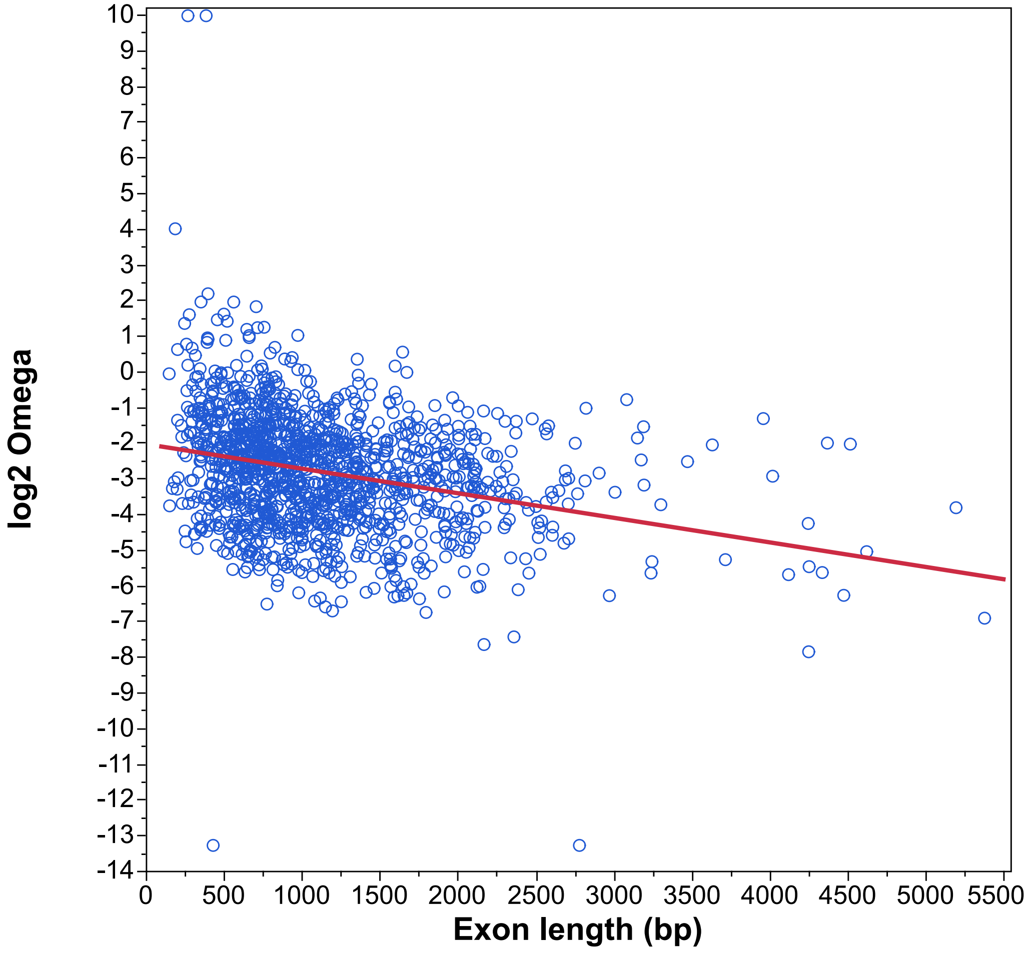

**Fig. S22** Relationship between $\log_2 \omega$ and exon length among single exon genes (1,372 loci) ($r^2 = 0.08$, $P < 0.001$).
